# The First 1,000 Days (1kD) Project - Collecting and Analyzing an Ultra-Dense Naturalistic Dataset of Human Baby Development

**DOI:** 10.64898/2026.03.19.712982

**Authors:** Hadas Raviv, Liat Hasenfratz, Kira Gousios, Marian Faryna, Ricky Beaty, Dean Johnson, Berlin Chen, Aja Altenhof, Brooke Ryan, Chandra A. Greenberg, Zhuoqiao Hong, Gal Assayag, Arkadii Tsyhanov, Valery Malakhov, Tal Rosenwein, Ofri Raviv, Casey Lew-Williams, Uri Hasson

## Abstract

Human development unfolds in continuous, multimodal environments across seconds, days, and years, yet most developmental datasets capture sparse, context-limited samples of everyday life. We introduce the First 1,000 Days (1kD) Project, an initiative designed to collect ultra-dense, longitudinal, child-centered data that capture developmental trajectories within their full ecological context. Fifteen U.S. homes with 17 infants were recorded 12-14 hours per day over a median of 944 days, yielding ∼1.18 million hours of raw audiovisual data. We present an end-to-end framework for large-scale longitudinal naturalistic measurement and a scalable analysis pipeline of the collected data. In a case study, we describe how we utilized our pipeline to isolate child-centered speech, resulting in the collection of 2,000 to 6,000 hours of transcribed speech for each infant. We demonstrate that dense sampling within the home environment reveals a stable, household-specific lexical structure, which sparse sampling methods consistently fail to capture. The 1kD project offers a blueprint for teams aiming to collect and analyze natural behavior at scale in real-world settings.

## Introduction

Human development is a dynamic, embodied, and multidimensional process. From birth, children inhabit environments saturated with continuously changing, multimodal input - acoustic, visual, and social - that unfolds across seconds, days, and years. Learning to move, perceive, interact, and use language requires integrating information across multiple time scales and modalities. To effectively characterize these processes, it is essential that measurements capture real life as it unfolds dynamically across multiple timescales, using naturalistic, extensive (ultra-dense) recordings. Doing so requires methods that capture children’s early development in ways that preserve everyday household environments, span temporal resolutions from seconds to years, and link a child’s behavior to the structure of the input they receive.

Until recently, such measurement requirements exceeded what was technically feasible. Assembling a large, continuous corpus that reflects everyday input and developing the tools to analyze it at scale seemed out of reach. Consequently, much of cognitive and developmental science has relied on methodologically controlled experiments, short-term observations, and carefully designed tasks, which have produced powerful insights into cognition, language, and social learning ^1–7^. Yet these approaches came at a cost - they strip away context, compress time, and reduce rich, multimodal phenomena into decontextualized units^8^. As a result, many core constructs have been defined in ways that fit what could be measured, and theories were built around the slices of behavior that laboratory methods could reliably capture.

Even within these constraints, the field persistently pushed observation closer to the flow of everyday life. Pioneering studies such as Hart and Risley’s weekly one-hour home recordings over 2.5 years represented an early attempt to document children’s natural learning environments^9^. This effort was followed by the *Speechome Project^10^* - an ambitious longitudinal study documenting one child’s first three years of life. Conducted before the consequential advances in deep learning, the project faced significant analytical challenges^11^. The resulting corpus remains somewhat underutilized and inaccessible for further research. Still, it reflected a conviction that predated the tools: that dense, naturalistic, fully contextualized measurement could fundamentally change what we are able to ask - and explain - about development^12^.

More recent advances have greatly expanded the reach of naturalistic measurement. Day-long audio recordings, ambient sensing, egocentric video, and wearable devices now make it possible to document children’s environments from their own perspective, capturing the multimodal richness of everyday experience^13–18^. Datasets such as SayCam^18^, BabyView^16^, SEEDLingS^19^, and the PLAY Project^20^ have broadened both the scope and diversity of coverage, substantially increasing the number of hours of naturalistic data available for analysis. Collectively, this growing body of work has established essential empirical and methodological foundations for ecologically valid developmental research. Yet all these datasets typically capture only thin slices of children’s developmental trajectory - hours to, at most, dozens of hours per household. As a result, researchers must infer what the unobserved majority of everyday life looks like, and conclusions inevitably rest on generalizing far beyond what is observable.

If no single child can be observed every day, all day, a common workaround is to build a large, naturalistic corpus by aggregating shorter recordings from many children and households^19,20^. Major community infrastructures - most notably Databrary^21^ - have been essential in making this strategy feasible by providing researchers with access to thousands of hours of richly annotated video from child participants across both experimental and naturalistic contexts. While this approach increases the total volume of data, it relies on an assumption that is rarely tested directly: that aggregating thin slices of recordings across many children is sufficient to characterize the richness and distinctiveness of children’s everyday experience over days and years. If, however, there is significant variability across families, hours, days, and contexts, then brief observations (“thin slices”) may fail to preserve the distinctive structure of any individual child’s experience. In that case, the result reflects an “average” household experience, even though no such experience actually materializes.

Here, we introduce the First 1000 Days (1kD) Project, a complementary initiative designed to collect ultra-dense, longitudinal, child-centered data that capture developmental trajectories within their full ecological context. The primary goal of this effort is to preserve, as densely as possible, the faithful experiences of individual children and to characterize in detail the statistical structure of their environments. This is a crucial step toward theories of language development that are grounded in the actual structure and variability of children’s everyday input.

The 1kD dataset documents the environmental inputs and behavioral outputs of 17 infants in 15 homes across the United States during the first approximately 1,000 days of life. This dataset utilizes multiple audio and video recording devices in each home, capturing 12 to 14 hours of footage per day. Running from February 2022 through May 2025, we recorded a median of 944 days per household (minimum 495, maximum 1194). Over the data collection period across all families, this effort yielded a corpus of approximately 1,180,000 hours of raw audio/visual recordings, from which thousands of hours of speech and activity per child have been isolated. By linking continuous, naturalistic measurements of a child’s input and output across diverse contexts, the 1kD dataset empowers integrative analyses of experience and behavior, providing an unparalleled opportunity to quantify, model, and understand development in its natural form.

Building the 1kD dataset required us to rethink, from the ground up, what it takes to do rigorous developmental science in real homes over long timescales. The paper is organized around the two core challenges: **(1)** collecting dense, ecologically valid data robustly and ethically, and **(2)** analyzing and quantifying the dense multidimensional data at scale. We describe the concrete framework we developed to meet these challenges and document the dataset’s structure and affordances for future developmental questions. Full implementation details are provided in the Methods and Supplementary Sections to support reuse, replication, and adaptation in other naturalistic studies.

We outline the frameworks developed for both data collection and data analysis. First, we highlight the most essential components of the ethical framework for recruiting and sustaining family participation over three years. Then, we describe the robust, secure, and scalable data collection pipeline we have developed. On the analysis side, we developed new infrastructure to process this naturalistic corpus at scale: 1) a human-centered annotation framework designed to evaluate the precision and recall of different artificial intelligence (AI) models in a naturalistic setup. 2) distributed computational systems leveraging dozens to hundreds of CPUs and GPUs to deploy these models efficiently across millions of audiovisual clips. The resulting analyses were stored in a centralized, continuously expanding database, designed to be enriched over time as new annotation tools and models become available.

Finally, we provide technical validation and an example case study by examining a central measurement problem in early language experience: what linguistic input do children have access to at home from early in life across different households, and how much of that input is household-specific? For each infant, we analysed thousands of hours of home speech processed through the pipeline. The analyses reveal both shared regularities and robust household-specific signatures in the linguistic environment. Crucially, we show that sparse or intermittent sampling does not preserve the distinctive structure of each child’s language input, highlighting the value of ultra-dense longitudinal recordings for developmental research.

This case study underscores the utility of the 1kD corpus for addressing fundamental questions in language acquisition and child development and provides a template for using the dataset and infrastructure to investigate a wide range of developmental phenomena. We hope that the 1kD dataset will spur the development of new computational tools and models for studying children’s early learning in naturalistic settings. More broadly, the methods and infrastructure established here offer a scalable framework for examining human behavior in real-world contexts across disciplines, including psychology, neuroscience, linguistics, and computer science.

## Results

### Family recruitment

With human subjects approval (Princeton University IRB, USA) to collect continuous naturalistic home recordings over three years, we recruited 15 families, each with a newborn infant or expecting a newborn. An ethical framework was developed before recruitment through regular workgroups with community stakeholders and experts (including the Institutional Review Board, General Counsel, and data security offices at Princeton University, as well as specialists in clinical and family psychology, maternal/infant health, and family financial planning). The ethical framework is informed in part by biobank governance models^22–24^, centered on fostering participants’ own control over the dataset: the right to delete any recordings they wish, the child’s right to delete their corpus when they are 18 years of age, and a clearly defined scope of use. Participant compensation was provided in the form of monthly contributions to a 529 college savings account for the focal child^25^.

Families could withdraw at any time. Recordings were held for two weeks before processing to allow families to request deletions; later requests were also reviewed and approved (101 requests across all 15 families were received; 297 hours total were deleted). Participants will be recontacted when the focal child turns 18, at which point they may request deletion of their corpus.

### Data structure

Fifteen families joined the project. The sample was recruited primarily in New Jersey, a highly diverse U.S. state. In aggregate, participating households reflected substantial variation in race/ethnicity, socioeconomic circumstances, family structure (including nuclear and multigenerational households), sexual orientation and gender identity, and daily routines and dynamics. Families lived in a range of housing contexts, including single-family homes and rental apartments, with variation in household size and physical space (from smaller to larger homes). Across households, the number of recording devices ranged from 4 to 14 (total = 132), and the number of recorded rooms ranged from 2 to 5 (57 rooms total).

The final sample comprised 17 children (9 female), reflecting the enrollment of two additional newborns born to participating families during the study period. Household composition varied: across families, 15 siblings participated (mean age 6.9 ± 2.2 years; 6 female). Several homes also included additional regular caregivers or relatives (e.g., nannies, grandparents, aunts/uncles; n=16; ages 22–62; 6 female) and pets (dogs/cats; n=18).

Data recording began in February 2022 and ended in May 2025. Recordings ended slightly earlier than planned due to funding constraints. By the end of the recording period, the median age of the 17 infants was 947 days (range: 288–1301 days). In total, approximately 1.18 million hours of video and audio information were recorded, with a range of 38,521–149,051 hours per household.

### Behavioral data collection

To complement the multi-modal home recordings, we collected longitudinal behavioral data, including demographics, caregiver-report questionnaires, and direct assessments spanning language, cognitive, motor, and socio-emotional development (see Methods for the full assessment battery and timeline).

The primary behavioral measure used in our analyses was the MacArthur-Bates Communicative Development Inventories (MB-CDIs^26^). Caregivers completed MB-CDI production reports monthly beginning at 8 months of age and continuing through each child’s participation. This sampling density is greater than is typical in the CDI literature, where deeply longitudinal repeated measurements are relatively scarce (as reflected in Wordbank’s available longitudinal corpora^27^. We administered the Words & Gestures form from 8–16 months and the Words & Sentences form thereafter. Between 8 and 16 months, children’s language comprehension was additionally assessed through an interview-based administration of the Words & Gestures form. The resulting high-frequency CDI series supports analysis of within-child (intra-individual) trajectories in observed language development, rather than relying only on between-child age trends.

### Data collection pipeline

In-home video recording is a well-established method in behavioral research^17^, but collecting ultra-dense, continuous data over 1,000 days poses unique challenges. The most significant among these are: **(1)** *Long-term robustness:* devices must function continuously and reliably with minimal maintenance for nearly three years; **(2)** *Minimal intrusiveness:* the devices should not disrupt family life in order to preserve natural interactions; **(3)** *Data security and privacy:* comprehensive and robust protection measures should be implemented to protect the sensitive data against unauthorized access, data breaches, and misuse; **(4)** *Scientific traceability:* the entire data lifecycle must be documented, from acquisition to transformation, ensuring visibility into any potential biases introduced during the process.

To address these demands, we developed a system comprising two tightly integrated components: (i) an in-home recording setup and (ii) a secure, cloud-based research environment on AWS for accumulating, storing, and processing the data.

### In-home recording setup

We utilized Wi-Fi cameras and far-field microphones to enable reliable, continuous recording. The main living areas in each home (e.g., family room, playroom, kitchen, dining room) were equipped with one to two cameras with embedded microphones and one additional high-quality microphone (Figure 1). Depending on the home’s size and layout, the total number of devices ranged from 4 to 14 (132 devices across households). All devices in each house streamed to a recording bridge installed, which collected recordings throughout the day and generated timestamps to support time synchronization across rooms and devices.

**Figure 1:**
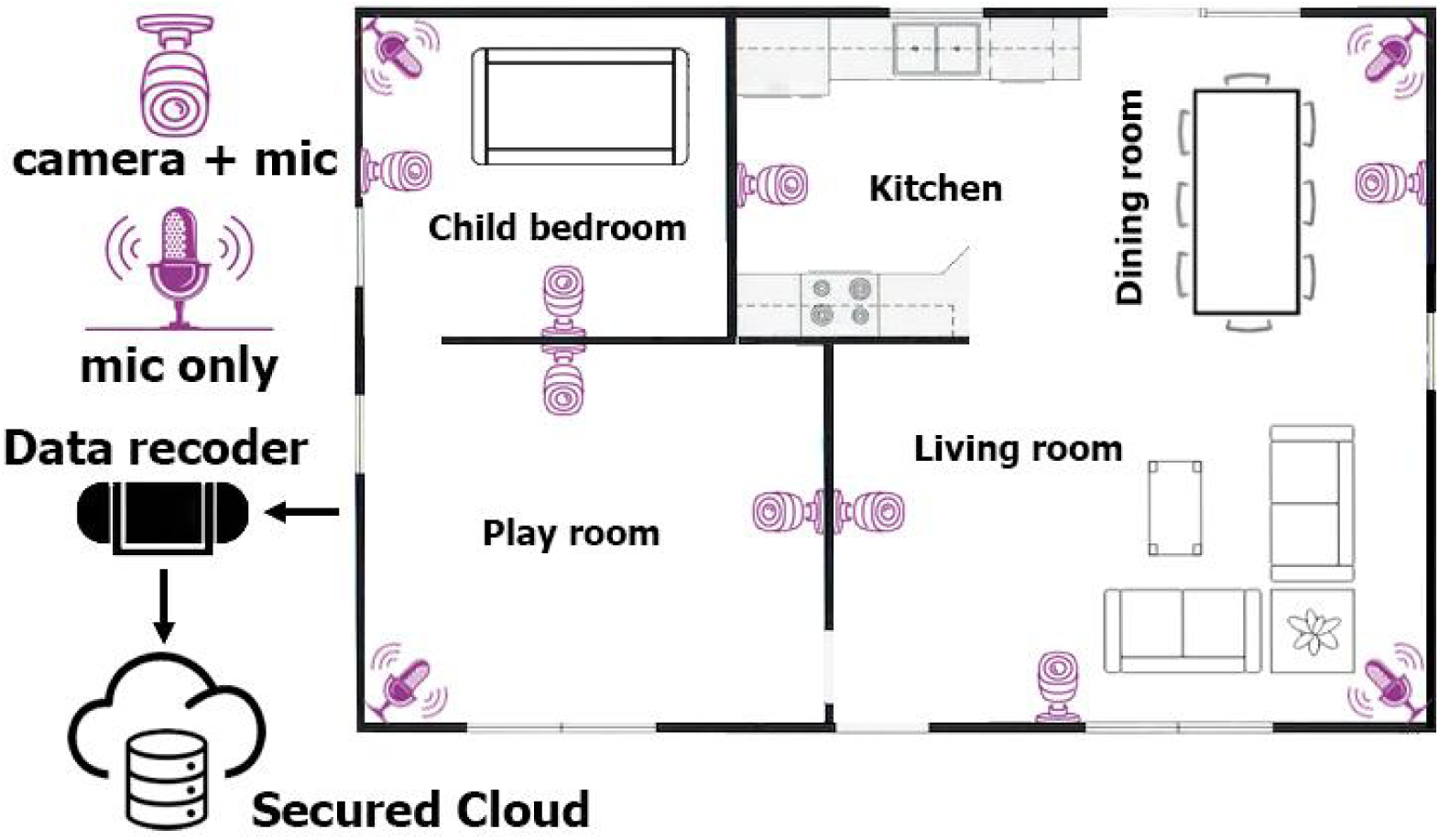
Setup of the recording system in family homes: the main living areas in each house (e.g., family room, playroom, kitchen, dining room) were equipped with wireless recording devices (audiovisual cameras and designated high-fidelity microphones). Data was transferred via a dedicated wireless network to a secure storage server and then to the AWS-based research environment.

Recordings were uploaded daily to an intermediate secure server and subsequently transferred to our secure AWS-based research environment. Full details are provided in the Methods section.

### Cloud-based download, processing, and storage pipeline

Every hour during the study period, our dedicated research system on AWS downloaded the daily recordings from the intermediate server to secure cloud storage. After the two-week cooling period, our data processing system was triggered to standardize the recordings. It split the set of audio–video and audio-only recordings into one-minute clips aligned to the clock (e.g., 14:03:00–14:03:59). This consistent, one-minute structure simplified searching, synchronizing rooms within a home, and analyzing data across different timescales. The one-minute clips were then transferred to long-term storage. The data download and processing systems operated continuously to maximize data collection, with automated retry modules that detected and recovered missing data to ensure completeness. Additionally, we maintained comprehensive metadata logging to support auditability and scientific provenance. Full technical specifications of the data-collection system are provided in the Methods section.

### Robustness and density of our recording system

The 1kD recording system proved to be robust over the 1,000-day collection period. Figure 2 summarizes daily recording density for each home, from the first to the last day of recording; a minute is considered “recorded” if any device captured a file during that time. Across all homes and **net** recording days, we achieved a median of 11.9 recorded hours per day, with daily averages ranging from 11.7 to 12.7 hours. In total, the dataset comprises ∼1.18 million hours of audiovisual recordings, equivalent to approximately 71 million one-minute files. Brief gaps occurred when individual devices failed, typically affecting specific rooms rather than entire homes, and temporary pauses occurred intermittently, most often when families were traveling. To increase coverage during periods when families were more likely to be at home, we extended the daily recording window from 12 to up to 14 hours for several families over the course of the study. Retention was high, with only two families relocated and ending participation after approximately 550 days.

**Figure 2.**
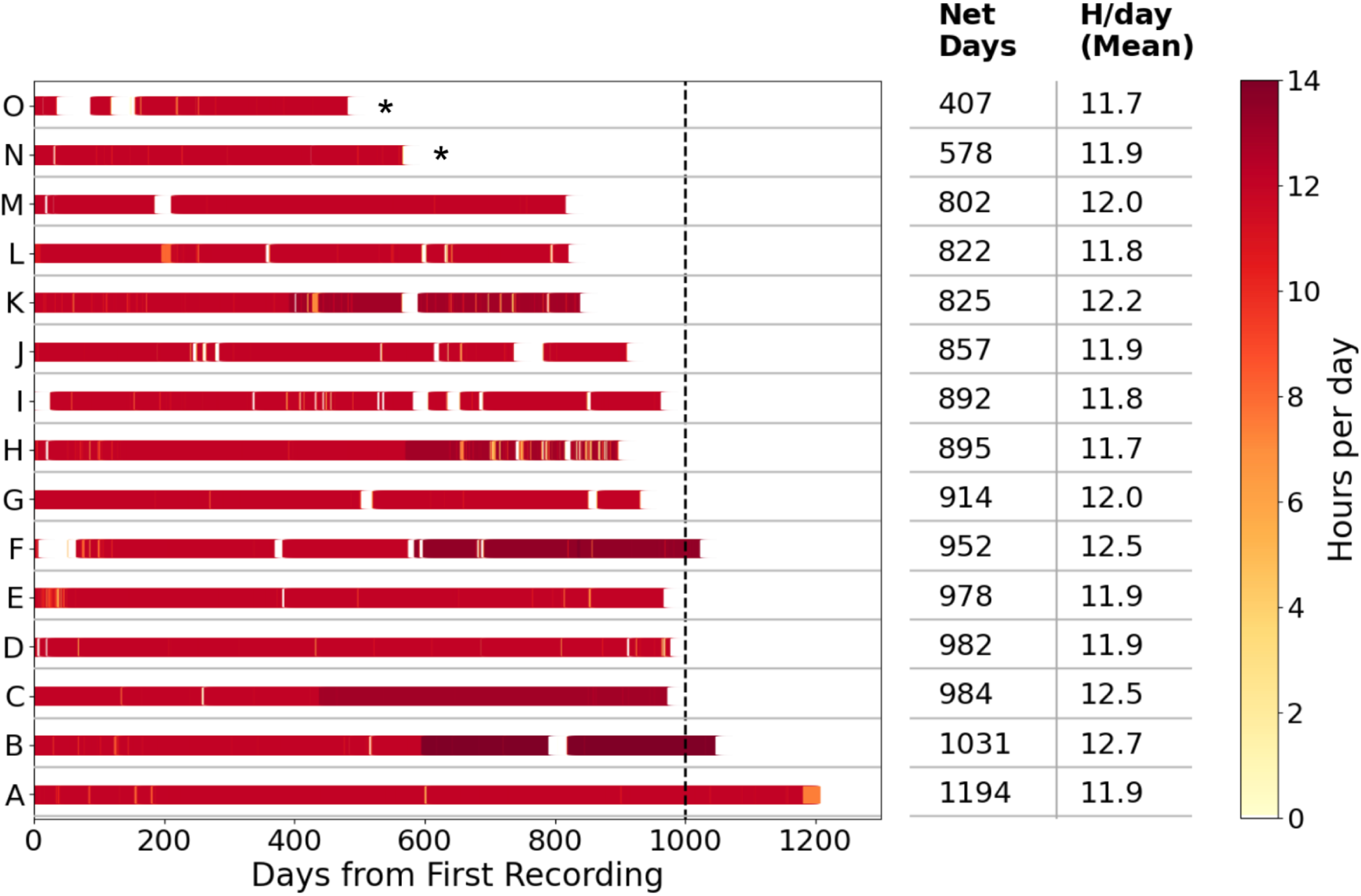
Daily recording density for each home from the first to the last day of recording: Each horizontal row represents a single house, with color indicating the number of recorded hours in the home per day, ranging from 0 (lightest) to 14 (darkest). A minute is considered “recorded” if at least one device from anywhere in the house was active during that time. This visualization highlights the system’s robustness and the variability in data coverage across homes and over time.

### The scalable data analysis pipeline

The 1kD dataset was designed to capture the richness of infants’ early experiences in ecological contexts, but the scientific value of such a corpus ultimately depends on the tools used to analyze it. Two dimensions of the data jointly create the central analytic challenge: the recordings are both highly naturalistic and unprecedented in scale.

### Naturalism

The recordings capture the complexity of everyday family life across rooms and across timescales from minutes to years. Audio frequently contains multiple simultaneous speech streams, sound sources, and background activities (e.g., cooking, dogs barking, media). Video is sometimes partially occluded, affected by shifting room layouts (e.g., holiday decorations), and includes everyday clutter. Household composition and activity patterns fluctuate over time. Critically, the child’s behavior changes rapidly over development: vocalizations progress from cries and coos to babble and words; posture, appearance, and motor behavior shift dramatically; and interaction patterns and routines become more complex. Recording quality also varies across devices and time, as spaces differ in layout, visibility, lighting, clutter, and acoustics. Overlapping cameras and microphones provide multi-view redundancy, introducing additional synchronization and reconciliation requirements.

Supplementary Figs. 1 and 2 illustrate this naturalistic complexity by showing, through random samples from two households, variability in where people are located and who is co-present across areas, as well as the frequent moment-to-moment co-occurrence of multiple sound sources.

### Scale

Given the vastness of the 1kD corpus, exhaustive manual annotation is infeasible, as it would require decades of human effort. Moreover, as will be detailed below, a substantial portion of the recordings capture low-value content, such as empty rooms, sleep, or periods when no one is present. Isolating behaviorally meaningful moments at this scale requires artificial intelligence (AI) tools that filter out uninformative segments and extract features that capture the structure and dynamics of everyday activity.

To enable scientific exploration while addressing these challenges, we developed a research-question-driven computational framework for scalable, automatic annotation of key behavioral and environmental dimensions. The core idea is to reduce naturalistic noise by integrating multiple signals when possible, while strategically running specialized models on targeted subsets of the data to make computation tractable at scale.

The computational framework is summarized in Figure 3. First, we define the behavioral or environmental dimensions of interest and the relevant time span (Figure 3-A), based on the motivating research question. Second, we identify candidate AI models for each target dimension and benchmark them against human-labeled reference samples using standard metrics (e.g., precision, recall, F1) and inter-rater reliability to provide an empirical upper bound on achievable performance (Figure 3-B). Model selection balances accuracy with computational cost, which is critical at this scale (e.g., a model that takes 1 second per file would require nearly 20,000 compute hours to process the whole corpus). Third, we deploy models at scale by packaging them as modular Docker containers and executing them on targeted subsets of the data via a cloud-based customized batch system that can launch tens to hundreds of CPU/GPU instances on demand (Figure 3-C). By restricting inference to relevant subsets (e.g., segments already identified as containing speech), compute-intensive models are reserved for cases where they add the most value. Finally, we integrate model outputs into a central feature table with full metadata (model version, parameters, runtime context) to support traceability and reproducibility (Figure 3-D). The 1kD feature table is modular and updatable, allowing new annotation layers to be added and prior layers to be refined as tools improve. Full implementation details are provided in the Methods section.

**Figure 3:**
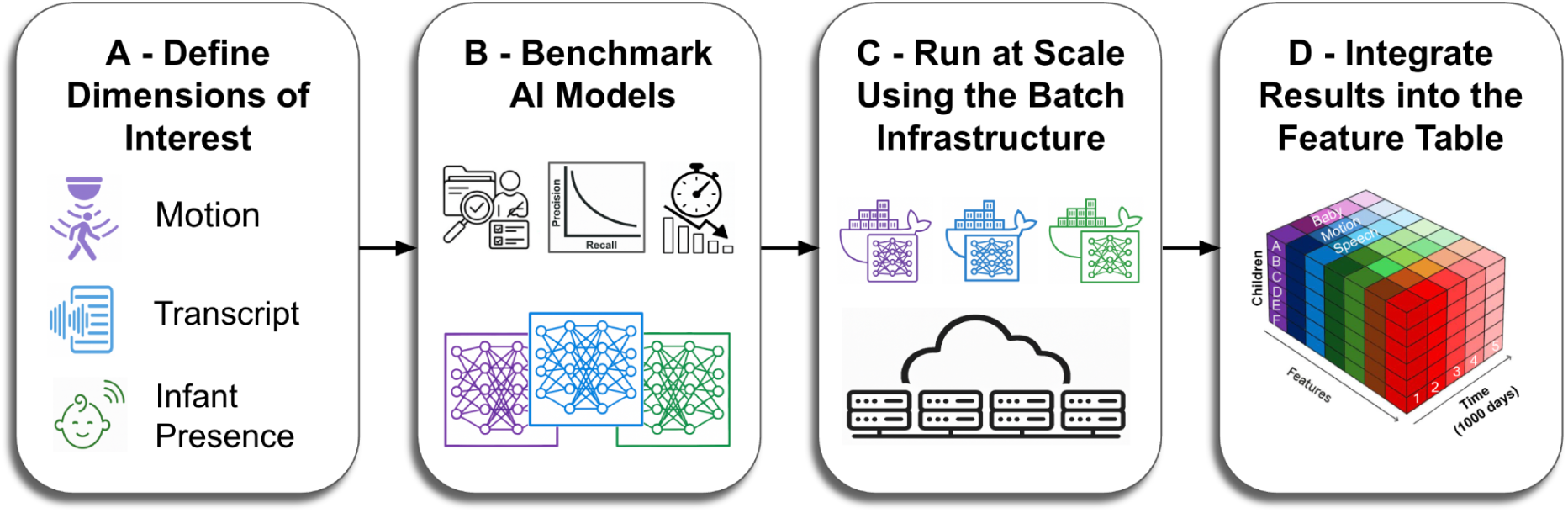
End-to-end framework for analyzing the 1kD dataset at scale. (A) Given a research question, researchers define the behavioral and environmental dimensions of interest (e.g., motion, speech transcript, infant presence). (B) AI models are evaluated for each dimension using manual annotation of a representative sample. This sample is used to assess each model’s performance and provides standardized metrics such as precision and recall. (C) Validated models are deployed at scale using tens to hundreds of CPUs and GPUs using a cloud-based batch infrastructure. (D) The resulting annotations are integrated into a unified, queryable feature table organized by child, feature type, and time, enabling flexible downstream analysis.

### Framework implementation case study: Isolating children’s language input

Many scientists have studied how early experiences influence development, spanning children’s cognitive, linguistic, and social-emotional lives. In the domain of language, significant efforts have been made to capture the nature of children’s language input as a source of variability in language learning^9,14,28–31^. Prior work was based on multi-hour to multi-day recordings, but scientists have not previously captured language input with the density and developmental continuity of the 1kD dataset. Here, we ask a foundational question: *how variable are the language environments across households, and how much sampling is required to characterize each one reliably?*

We use this question as a case study to demonstrate the capabilities of our processing framework (Figure 3) and to provide a basis for evaluating existing language input estimates and testing theoretical assumptions about children’s everyday experiences.

Characterizing the language environment in a longitudinal, multi-room, multi-device recording setup requires fusing complementary cues: audio to identify when speech occurs, and video to determine when and where the infant is present. We implemented this fusion in a room-level, minute-resolved pipeline that produces probabilistic, child-centered aggregate annotations and an associated speech corpus. To ensure computational efficiency at scale, the pipeline was implemented hierarchically: initially, computationally lightweight methods were used to screen the data, with higher-cost models applied only to targeted subsets. Below, we summarize the pipeline shown in Figure 4; full implementation details of each algorithm are provided in the Methods section.

**Figure 4.**
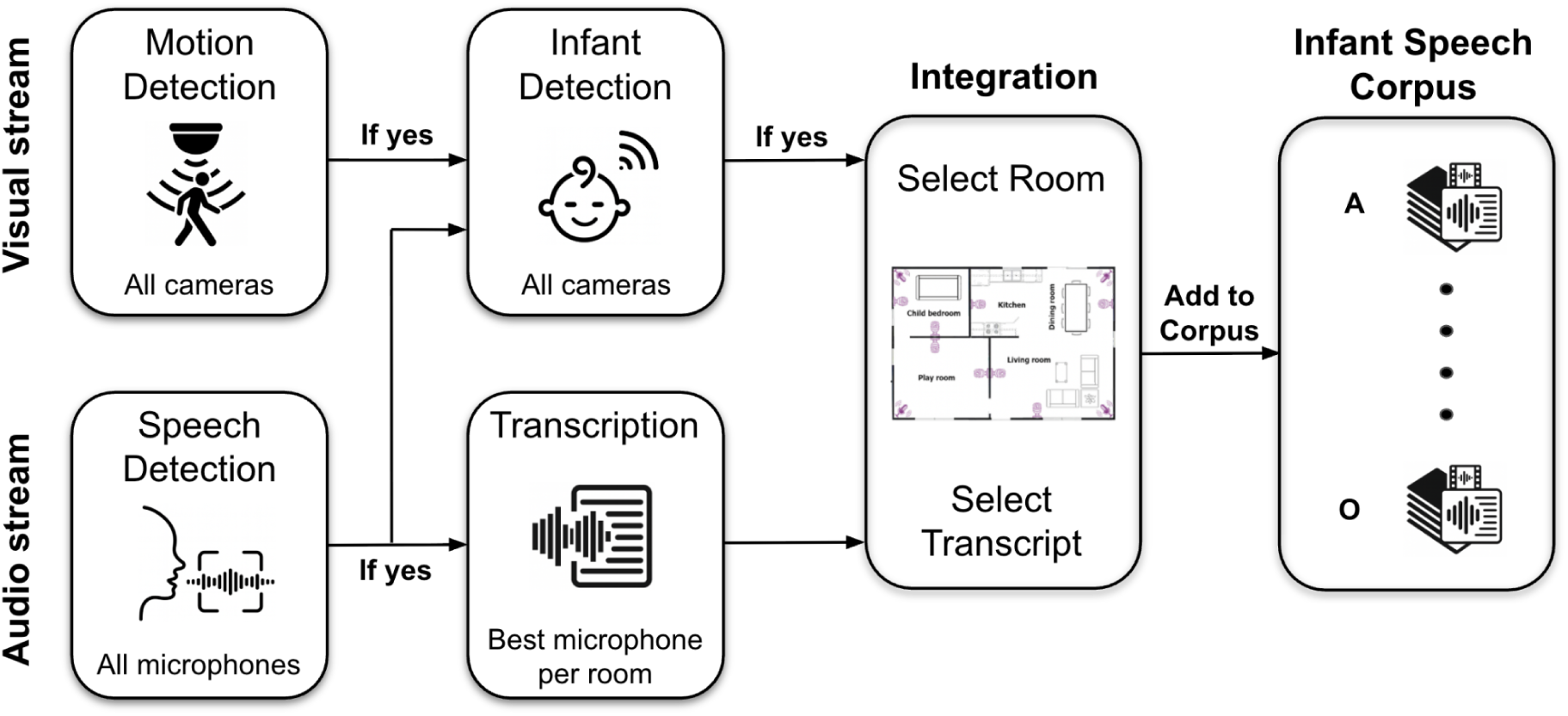
Overview of the audio–visual pipeline used to generate infant-centered speech segments. In the audio stream, a speech detector first identifies minutes containing speech. For those minutes, the highest-energy microphone recording is selected and transcribed. In parallel, the video stream is processed using motion detection. Minutes that contain both motion and speech are further annotated for the presence of the infant and other individuals. For each minute, an infant presence score is computed for each room; if any room exceeds a predefined threshold, the transcript associated with the highest-scoring room is added to the child-specific corpus.

### Visual processing stream

For each recorded minute, we first applied a lightweight motion-detection algorithm to all one-minute video segments. We then aggregated motion signals across cameras within a given room to determine whether any activity occurred in that space during each minute. This stage was tuned for high recall, capturing movement from either people or pets. To further focus processing on infant-relevant moments, we intersected the motion annotations with the audio-based speech detector. Subsequent visual inference was run only for minutes in which (i) motion was detected, and speech was present, or (ii) speech alone exceeded a conservative threshold (above 8 seconds of speech per minute). For these selected minutes, we applied a multimodal model to detect the infant and other household members (e.g., adult female, adult male, older sibling). To date, we have applied infant detection to eight families. Model outputs were aggregated across cameras within each room to yield, for each minute, an inferred probability of the presence of each person in each room.

### Audio Processing Stream

For each recorded minute, we used a wav2vec-based speech detector^32^ to identify minutes containing speech; minutes with detected speech were carried forward for processing. We then applied a denoising model to suppress background noise and better isolate vocal content^33^, and computed overall speech energy for each minute. For each room and minute, we selected the recording with the highest signal energy as the representative speech signal for that space. These selected recordings were transcribed with a state-of-the-art automatic speech recognition model, WhisperX^34^, yielding one transcript per room per minute. This audio pipeline was applied to all families in the dataset.

### Integration

At the final step, we combined the visual and audio outputs using temporal continuity constraints to smooth and fuse minute-level predictions across rooms. This integration produced, for each minute and each region of the house, an *infant presence detection score* (ranging from 0 to 1) representing the likelihood of infant presence. To construct the child’s language corpus, we first identified minutes in which at least one room’s infant presence score exceeded a predefined threshold. For those minutes, we selected the transcript associated with the *highest-scoring room* to represent the child’s language environment. As discussed below, the resulting child-centered speech corpus can be tuned to different research goals: lower thresholds on the infant presence score prioritize recall, capturing a broader range of contexts (at the cost of increased noise), whereas higher thresholds prioritize precision by including only moments with stronger evidence of infant presence.

### Pipeline performance metrics

To evaluate end-to-end pipeline performance (Figure 4) across the eight thoroughly analyzed families, we randomly sampled 100 one-minute segments per household from the full recording period. Each sampled minute was independently labeled by human annotators for the presence of (i) the infant, (ii) another child, and (iii) speech. To contextualize the model’s performance relative to human agreement (machine-human), we estimated inter-rater (human-human) reliability by computing Cohen’s Kappa on 20 minutes randomly sampled across households, annotated by two independent raters per category. Using the annotation labels, we computed the mean precision, recall, and F1 for each category (infant presence score threshold = 0.1). Because prevalence varied substantially across households (≈20–55% of minutes positive for infant presence), we also report mean precision lift, defined for each household as observed precision divided by the chance-level precision implied by that household’s prevalence. Table 1 reports performance for (i) infant presence, (ii) any child presence (infant or other child), (iii) speech presence, and (iv) the intersection of child presence and speech. All reported results are averages across households.

**Table 1.**
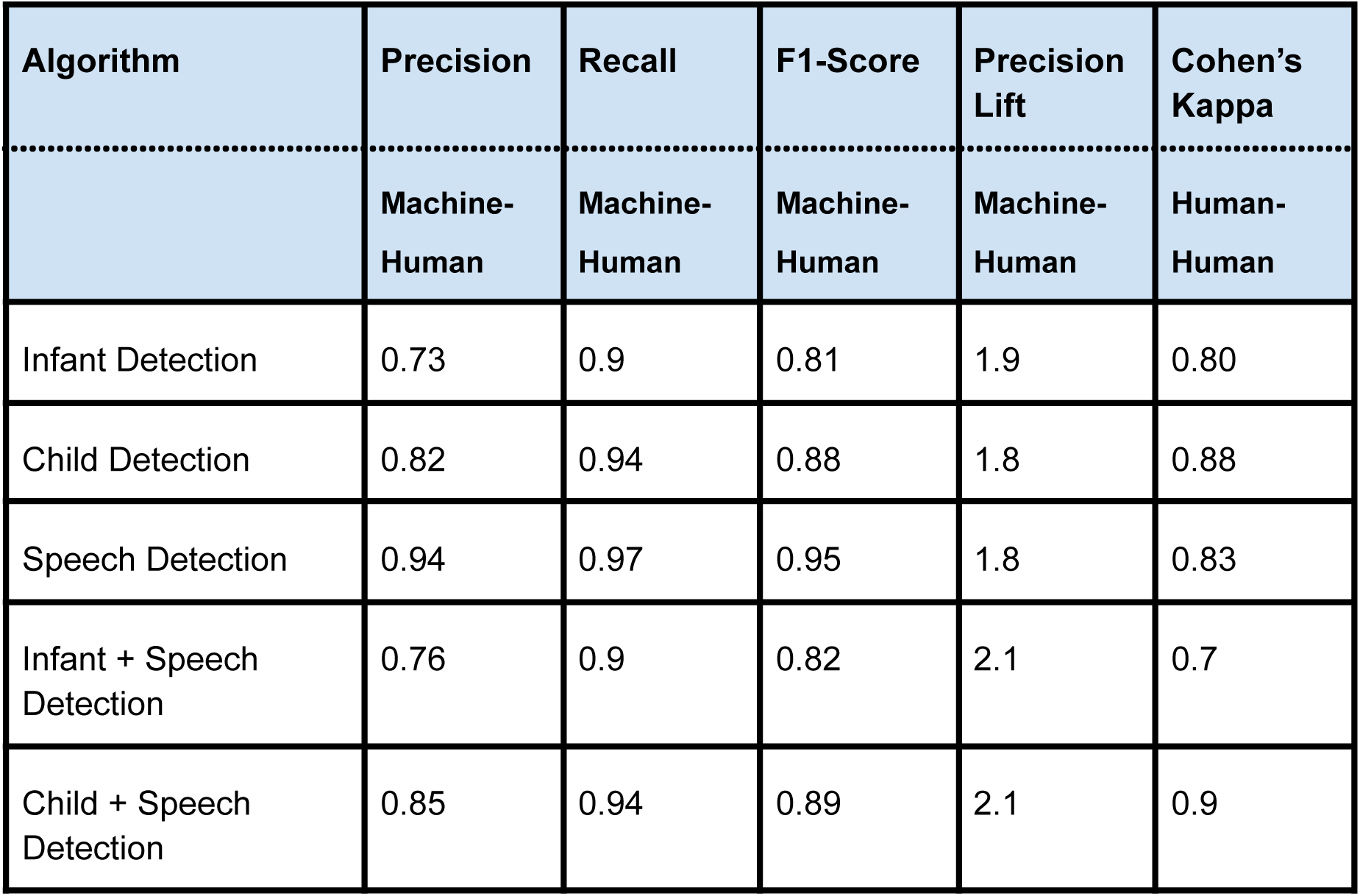
End-to-end detection performance of the home pipeline averaged across eight families. We report precision, recall, F1, Precision Lift, and Cohen’s Kappa scores for infant presence, any child presence (infant or other child), speech presence, and the intersection of child and speech presence. Metrics are computed from 100 randomly sampled minutes per family. The infant presence score threshold was set to 0.1, and speech presence was defined as> 1 word in the WhisperX transcript.

The F1 score was high (above 80%) for all levels of analysis (Table 1). The speech-detection pipeline performed exceptionally well, achieving very high precision (94%) and recall (97%) in determining whether speech is present in the home. Infant-detection performance was strong (F1 of 81%, precision 73%, recall 90%). It was further improved when considering the presence of any child (F1 of 88%, precision 82%, recall 94%) and when intersecting with the speech (F1 of 82%, precision 76%, recall 90%). Importantly, the models’ performance was comparable to that of human raters, as measured by inter-rater reliability (Cohen’s kappa; Table 1).

In the Supplementary section, we report the full performance of the algorithms used in the pipeline. Across households, all algorithms achieve high overall precision and recall. Our scientific analyses are based on the final integrated pipeline output rather than on any single component in isolation. We also quantify child- and speech-detection performance as a function of the infant- and child-presence score thresholds, showing that the precision - recall tradeoff can be controlled by setting different values for this parameter.

### Transcription quality

To evaluate transcription quality, we manually annotated 183 one-minute audio samples randomly selected from the eight thoroughly analyzed homes. Each sample was split into two 30-second intervals for transcription, yielding 366 segments in total. Annotation guidelines were developed iteratively to accommodate the 1kD multi-room recording setup and are described in the Supplementary Materials. Annotators were native speakers who trained on as many examples as necessary to pass evaluation and completed all annotations using headphones.

The recordings span the full range of everyday acoustic variability: diverse sound sources, speakers moving between rooms, overlapping speech, and large minute-to-minute fluctuations in speech rate. For both corpus construction and evaluation, we exclude samples that are unlikely to contribute meaningful linguistic content using two criteria. First, we remove low-quality transcripts with a high *compression ratio*, defined as the original text size divided by the size after zlib compression (both in bytes). High values indicate unusually repetitive, low-information outputs that are often associated with ASR hallucinations^35^. These account for 7.9% of the evaluated segments. Second, we exclude very short transcripts (<= 5 words), which typically reflect extremely brief speech or faint/background speech that cannot be reliably transcribed. These account for <1% of the total words in the final transcription-based dataset.

Overall, human and model outputs were closely aligned. Word counts were highly correlated (r = 0.84 and 0.83 before and after short transcripts filtering, respectively), indicating that the model tracked overall speech quantity well. Accuracy metrics are also comparable across human-model and human-human comparisons. Restricting to segments containing at least five words in both the model and human transcripts, we obtain WER = 0.51, MER = 0.46, precision = 0.74, and recall = 0.59 for human–model comparisons. Under the same length criterion, human–human comparisons yield WER = 0.49, MER = 0.47, precision = 0.75, and recall = 0.55. This similarity likely reflects the prevalence of background or partially audible speech in naturalistic recordings, which makes it difficult - even for humans- to decide what should be transcribed. Overall, these values fall within the expected range for naturalistic speech transcription, especially in noisy, natural conditions beyond laboratory-controlled settings^36^.

Finally, a small set of cases in which WhisperX produced relatively long transcripts while human annotators transcribed few words can be interpreted as potential hallucinations; these were rare (∼1% of segments). However, manual inspection suggested that in roughly half of these cases, a new annotator judged the WhisperX transcript more plausible than the human annotation, in alignment with the human-human comparison scores. Taken together, these findings indicate that the corpus is dominated by segments with substantial speech content and high transcription quality, with WhisperX performing particularly well relative to human transcription.

### Total number of hours containing speech

We applied the **audio-stream** pipeline to all families. As shown in Extended Data Fig. 4, the total recorded hours varied by family, and the amount of isolated speech (regardless of child presence) ranged from 2,370 to 10,983 hours per family (median = 5,655). Differences in the overall proportion of speech hours across families reflect variation in time spent at home and in the intensity of vocal activity.

In addition, we intersected available speech at home with the infant’s presence in **eight** families. Table 2 reports the total hours collected for these families, including unique speech hours, unique infant hours, and unique infant+speech hours. The dataset contains thousands of hours of infant-and speech-related moments per child (ranging from ∼1,500 to ∼6,200 hours) and is two orders of magnitude larger than any previously collected dataset^16,18^. We note that the infant presence score threshold used here is 0.1; lowering it would increase the estimated number of hours by several hundred per infant. Adding any child presence criteria will further increase the number.

**Table 2.**
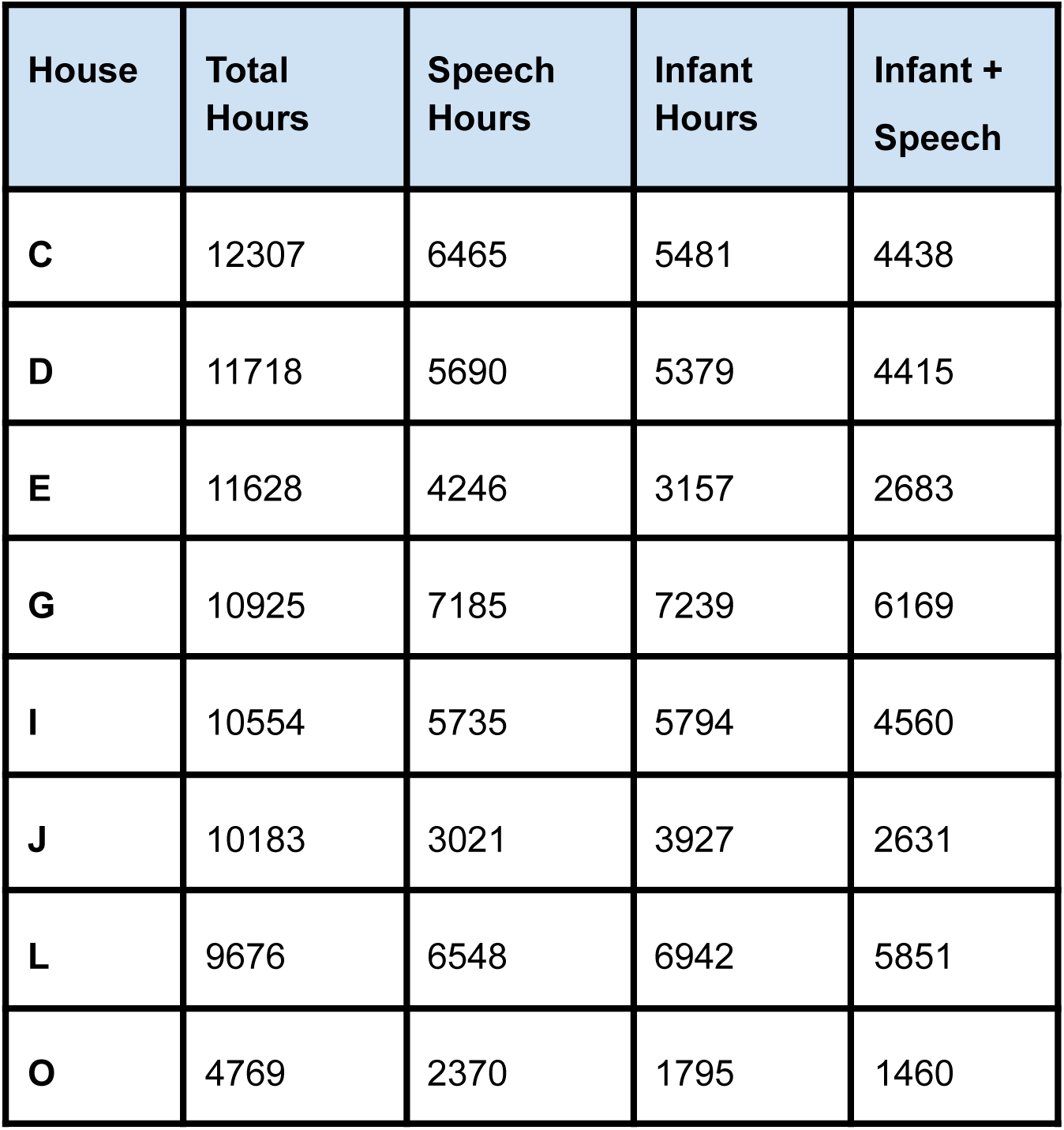
Total recording hours, unique speech hours, unique infant hours, and unique infant+speech hours for fully processed families in the 1kD dataset.

| House | Total Hours | Speech Hours | Infant Hours | Infant + Speech |
| --- | --- | --- | --- | --- |
| C | 12307 | 6465 | 5481 | 4438 |
| D | 11718 | 5690 | 5379 | 4415 |
| E | 11628 | 4246 | 3157 | 2683 |
| G | 10925 | 7185 | 7239 | 6169 |
| I | 10554 | 5735 | 5794 | 4560 |
| J | 10183 | 3021 | 3927 | 2631 |
| L | 9676 | 6548 | 6942 | 5851 |
| O | 4769 | 2370 | 1795 | 1460 |

### Linguistic properties of natural home environments

The extensive 1kD recordings provide indispensable new insights into core questions in developmental science. As a case study, we focus on early language experience: the quantity and richness of speech children encounter in their natural environments from very early in life. A detailed linguistic analysis will be presented elsewhere (Raviv et al., in prep.); here, we report findings that motivate this paper and directly address two open questions: (1) whether an “average” household is a faithful abstraction of real home language environments, and (2) whether thin-slice sampling can characterize household language properties without substantial information loss. As a baseline, we compare our in-home language corpora with a widely used aggregated language resource, CHILDES^37^, a shared, open repository of child language corpora, comprising many datasets contributed by research groups worldwide.

For each language corpus, we constructed a vocabulary that includes all words produced in that environment, along with their frequencies. As described in the Methods section, all corpora were automatically segmented into sentences and tokens, and each token was assigned a lemma and part-of-speech (POS) tag. Vocabulary items were defined as unique lemmas. We also conducted an analysis based on lemma-POS combinations, which yielded the same general patterns. Vocabulary item (word) frequency was calculated as the total number of occurrences of that item, normalized by the total number of tokens in the corpus. To reduce noise from extremely rare words, analyses were restricted to items that occurred at least 10 times in each analysed dataset.

First, we analysed the frequency distributions of each individual vocabulary. We fitted the rank–frequency distributions of word types in each corpus using the Zipf–Mandelbrot equation^38–40^. As shown in Figure 5-A, the fitted distributions closely matched the empirical data. This close correspondence was further reflected in the Pearson correlation between observed and model-predicted frequencies, which was consistently very high across corpora (*r* ≈ 0.99).

**Figure 5.**
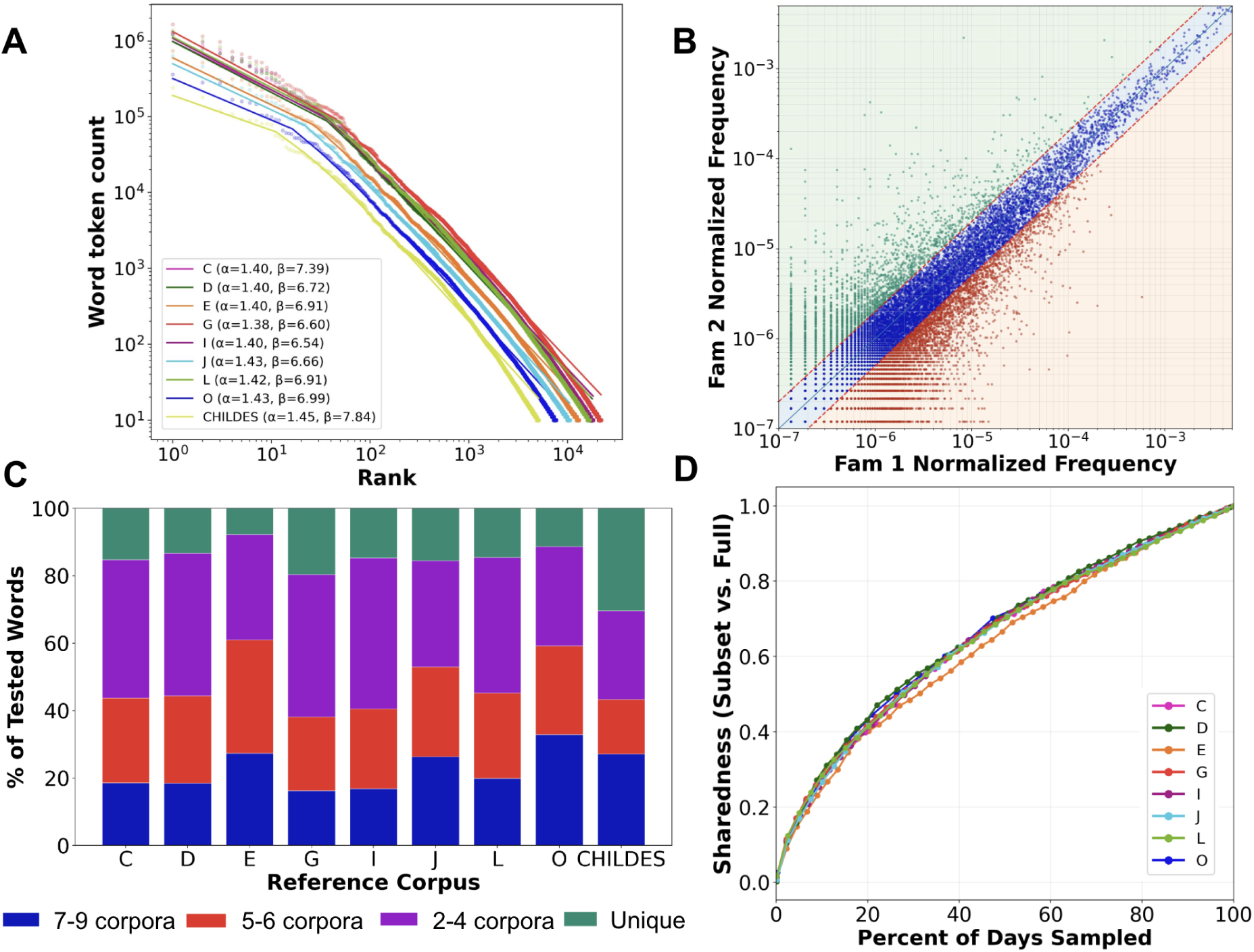
Language properties across households. (A) Log-log rank–token count curves for eight 1kD households and CHILDES show highly similar distributions, with closely comparable fitted parameters. (B) Example pairwise comparison of normalized word frequencies between two households (log-log scale). The blue region marks the fixed twofold band in which frequencies are considered consistent across households, whereas green and red indicate household-specific deviations. (C) Sharedness analysis. For each reference corpus, stacked bars show the fraction of words whose frequencies matched, within a fixed twofold band, in 1 corpus only (unique) or across 2–4, 5–6, or 7–9 corpora overall, indicating partial overlap and household-specific lexical signatures. (D) Within-household subsampling analysis: vocabulary sharedness (sampled subset vs. full corpus) increases with the number of sampled days, but converges slowly and similarly across households.

Together, these results indicate that the Zipf–Mandelbrot model provides an excellent characterization of the frequency structure of each household vocabulary and that this structure is a shared property of language across household environments. Importantly, the fitted parameters were comparable between individual households and CHILDES, suggesting that everyday communicative language has a similar overall frequency structure across these environments.

Although households showed similar overall frequency structure, the frequencies of individual lexical items varied substantially. Figure 5-B presents a representative comparison between two households: each point corresponds to a word, and the x- and y-axes represent its normalized frequency in the two corpora. Many words cluster within the fixed twofold band around the diagonal (blue), reflecting similar frequencies across households. These items can be considered part of a shared vocabulary, both in terms of lexical overlap and comparable frequency in naturalistic language use. In contrast, words outside this region (red and green) exhibited more household-specific frequency patterns, suggesting that individual households are characterized by partially unique lexical profiles. These findings indicate that an “average household” representation is insufficient to capture the structure of real language environments. Each household appears to have its own lexical signature, defined by words with distinctive frequency profiles, and averaging across households would obscure this specificity, yielding a flattened distribution that does not accurately represent any individual household.

To evaluate whether any of the tested corpora could represent an “average” household, we conducted a reference-based comparison across nine corpora: eight 1kD households and CHILDES. For each reference corpus, we compared all remaining 1kD corpora at the level of individual words. This yielded seven comparison corpora when a 1kD household served as the reference, and eight when CHILDES served as the reference. Word probabilities were estimated using additive smoothing, and for each word we computed the ratio of its probability in the comparison corpus to that in the reference corpus (see Methods). A word was considered shared if this ratio fell within a fixed twofold band, a deliberately permissive criterion designed to capture broad similarity in naturalistic frequency estimates. For each word in the reference vocabulary, we counted the number of comparison corpora in which it was shared with the reference, and then added the reference corpus itself so that the final count reflected the total number of corpora sharing that word. Each reference corpus was summarized by the proportion of words that were unique to the reference corpus or shared across 2–4, 5–6, or 7–9 corpora.

Lexical profiles varied greatly among families, with substantial differences in word frequency across households. Figure 5-C shows that, across reference corpora, the largest proportion of words (∼52% on average) was either unique to a single family or shared by only 2-4 corpora. This indicates that frequency agreement is typically observed across only a limited subset of households, rather than consistently across all corpora. Most of the remaining words were shared more broadly, with about 25% occurring in 5-6 corpora and about 23% in 7-9 corpora. CHILDES showed a comparatively larger unique segment, consistent with greater divergence in item-level frequency structure, potentially because it aggregates many distinct lexical profiles into a single distribution. Overall, item-level frequency profiles are not captured by an “average household”: many words generalize only partially across homes, and averaging yields a flattened distribution that is not observed in individual households. We observed similar patterns when we repeated the analysis for the major lexical classes of verbs and nouns (Supplementary Fig. 4-A,B). However, verbs were more consistently shared across households than nouns. This suggests that nouns are more closely tied to household-specific language relating to people, objects, and routines.

Finally, we assessed whether thin-slice sampling within a household can recover each household’s unique language distribution. For each household, we treated the full corpus as the reference, randomly permuted recording days, and constructed nested subsets consisting of the first *k* days in the permutation, increasing *k* stepwise. For each subset, we computed the word-frequency distribution over the fixed vocabulary defined above and compared it with the full corpus using the same fixed twofold band, again treating this as a permissive criterion for broad similarity. Figure 5-D plots the fraction of vocabulary words whose probabilities remained within this band relative to the full corpus as a function of sampling depth, expressed as the fraction of recorded days sampled (median = 805 days). Match rates increased with sample size, but only gradually. This slow convergence likely reflects the Zipf-like, heavy-tailed structure of lexical distributions, because closely approximating the full corpus requires accumulating evidence for many low-frequency words in the long tail. Notably, the match-rate curves were very similar across households, consistent with the broader finding that households share a common rank-frequency structure while maintaining distinct lexical profiles. These results highlight the value of 1kD ultra-dense, long-term recordings for characterizing children’s idiosyncratic vocabulary in real-life language environments.

## Discussion

Progress in our understanding of child development is driven by theory, methodological innovation, and debate, yet it remains profoundly constrained by the data we can collect. This research project aims to expand the empirical and theoretical basis of developmental research by collecting a unique, individual-level dataset spanning children’s first 1,000 days – one that is both ecologically valid and rich in developmental detail. To achieve this goal, we built a robust, longitudinal data-collection infrastructure and a scalable analysis pipeline for analyzing large-scale naturalistic recordings. Using these systems, we followed 17 children from 15 homes during approximately the first 1,000 days of their lives (median = 945). We then preprocessed and analyzed the collected audiovisual recordings to detect each infant in the videos and transcribe speech occurring in each infant’s vicinity. The resulting corpus provides an unprecedented record of home language input: approximately 2,000–6,000 hours of (automatically) transcribed speech per infant. Beyond the data itself, the 1kD project offers a blueprint for teams aiming to collect and analyze natural behavior at scale in real-world settings.

Our language analyses revealed a combination of shared statistical structure and pronounced household-specificity in children’s everyday language environments. Although lexical distributions across households were consistently well characterized by the same Zipf-Mandelbrot form, the frequencies of individual words differed substantially, indicating that no single corpus can serve as an adequate “average” household. Moreover, within-household subsampling showed that these household-specific lexical profiles can be recovered only through dense, long-term sampling, and that small samples do not accurately capture the full distribution of words present in naturalistic input.

### Framework for analyzing real-life corpora

We built a novel analysis pipeline that analyzes millions of audio-visual recordings at scale using state-of-the-art AI models. The system can deploy tens to hundreds of GPUs on demand and aggregate model outputs in a scalable database. The analysis pipeline is model-agnostic: it supports rapid experimentation with alternative models and, as models improve, enables easy deployment of new models and re-analysis of the full dataset at scale.

Crucially, by utilizing both human-human and human-model benchmarks, we developed a validation paradigm to estimate each model’s precision and recall *before* large-scale deployment. The evaluation numbers for the final aggregated annotation scores enable flexible selection of data subsets depending on the downstream scientific question: lower infant presence detection score thresholds prioritize completeness to build a rich, nuanced corpus, while more conservative thresholds allow the estimation of refined metrics needed to answer subtle questions.

An additional key contribution is the 1kD feature table, a modular, multi-layer repository designed to accumulate annotations across multiple dimensions in a flexible, ever-expanding manner. This table allows researchers to build on previous analyses to evaluate and select data in an interactive, context-sensitive way, while enriching the narrative with each analysis. Thus, each new dimension can interact with existing ones, significantly expanding the dataset’s scientific utility.

### On the uniqueness and importance of ultra-dense sampling

The 1kD dataset is unusual in both scale and scientific value. Each child contributes thousands of hours of infant presence, speech, and their intersection^16,18^. The ultra-dense recordings of individual children’s lives enrich developmental research by allowing empirical investigations of how unique contexts and within-child variability shape each child’s distinct developmental path. Our analyses show that aggregating short recordings across households is not equivalent to dense within-home sampling: thin-slice recordings often fail to recover the speech distribution of individual households. Aggregating daily recordings across families can’t uncover family-specific structure beyond a shared high-frequency core.

Dense measurement thus does more than increase volume-it allows us to *distinguish what is truly shared across homes from what is systematically household-specific*, and to test when aggregation is informative versus when it obscures meaningful structure. The practical sampling question depends on the phenomenon of interest. Capturing the relationship between the full set of words a child is exposed to over a developmentally relevant period and the subset that is ultimately learned requires long, intensive recordings. By contrast, analyses restricted to the most frequent words would require less sampling. This is precisely the aim of the 1kD project: to explicitly measure the relationship between developmental phenomena of interest and their ecologically valid input distributions.

The implications extend beyond language development: sparse sampling tends to privilege high-frequency aspects of experience, potentially missing less frequent but developmentally significant patterns. Many theories of learning emphasize the importance of variability, contingency, and rare but privileged events^41–47^ - precisely the kinds of experiences that are often overlooked without dense, context-rich observation.

### On the scientific importance of the dataset

The 1kD dataset’s density and multimodality enable the integration of multiple environmental factors - including speaker identity, speech style, and socio-temporal context - with detailed measures of infants’ developing behavior. This makes it possible to investigate how the fine-grained, multimodal statistical structure of input (across seconds, minutes, hours, days, weeks, months, and years) iteratively shapes a child’s unique developmental path.

#### Dynamic linguistic input and word learning

A primary frontier enabled by 1kD is the investigation of how context-dependent variability in linguistic input shapes individual vocabulary growth. Traditional measures often rely on aggregate word counts, yet our data reveals that each child is exposed to a unique ‘word diet’ shaped by household-specific patterns of nouns, verbs, and modifiers. By linking dense transcripts with longitudinal behavioral assessments such as the CDI^26^ and QUILS^48^, researchers can now assess how the "long tail" of infrequent words and the temporal structure of everyday life - whether bursty, periodic, or context-specific - influence the robustness of word learning.

#### Social interaction and caregiver dynamics

The multimodal nature of the 1kD dataset enables the study of the social and environmental signals that shape learning. Using modern multimodal (audiovisual) language models, we aim to systematically analyze key characteristics of the input, such as the ratio of child-directed speech to adult-directed speech in different contexts.

Our goal is to determine whether consistent features of this exposure are linked to developmental outcomes of individual children in real-life contexts. Beyond language, additional dimensions of interaction, such as affective tone, contingent responsiveness, proximity and co-presence, and patterns of activity context (media use, play, meals), offer a path to modeling how everyday dynamics support (or constrain) developmental trajectories.

#### Agentic modeling of human development

The scale and statistical richness of the 1kD dataset provide a foundation for a new class of "child-centric" agentic models. Modeling development in biologically and cognitively plausible ways is an active research field^49–51^. Progress, however, is bottlenecked by static, text-heavy corpora designed for engineering benchmarks and by datasets assembled from thin, fragmented samples aggregated across infants and households. In contrast, human development - and an artificial agent modelling it - requires continuous learning from noisy, temporally structured, and context-dependent inputs. By capturing longitudinal trajectories, the 1kD corpus allows us to move beyond building models trained on limited data to constructing mechanistic models that learn, like children, from ecological longitudinal inputs embedded in rich social environments. These shifts can uncover essential computational processes for processing the complexity and unpredictability of real-world environments, potentially revealing new hypotheses about human development.

In conclusion, the 1kD project presents a new framework for capturing and analyzing the complexities of early childhood on a large scale among a diverse group of infants in the U.S. By combining ultra-dense longitudinal data with transparent analytics and flexible features, we facilitate the exploration of previously untestable questions. This includes examining multidimensionality, individual variability, and the influence of environmental and social factors. We envision 1kD as a catalyst for a multidisciplinary effort that spans psychology, linguistics, neuroscience, and computer science. Our goal is to establish shared benchmarks and develop cognitively grounded models that continuously learn from natural experiences, much like how a developing infant learns about their multifaceted world, moment by moment.

## Methods

### Family recruitment and enrollment

Families were recruited via leaflets in daycare centers and announcements on Facebook. Before enrolling, interested families first watched a short video introduction to dense naturalistic recording (“The Birth of a Word”, TED Talk by Deb Roy^52^). Then they participated in a series of meetings with the research team, specifically with the project coordinator, our research assistant, and a faculty member overseeing recruitment. Across these meetings, we discussed the study’s scientific goals and rationale, walked through the recording setup and day-to-day implications of participation, and reviewed privacy/security safeguards and data governance in detail, with ample time for questions. We encouraged each potential participating family to provide input about how the study would work.

All meetings took place prior to consent. As part of the enrollment and participant-support procedures, families were also introduced to an independent clinical psychologist (not part of the study team), who served as an additional point of contact throughout participation.

### Data acquisition system

Our data-collection system comprised two coordinated parts: (1) behavioral data collection and (2) an automated technical pipeline integrating three components: (i) an in-home recording setup, (ii) a cloud-based data-transfer service, and (iii) a cloud-based system for data normalization and processing. Audio-video data recorded in the home were uploaded daily to a secure intermediate server for temporary storage and then transferred to, and processed within, our secure AWS-based research environment (components ii and iii).

### Behavioral data collection

We collected longitudinal behavioral and developmental data including demographic information (for example, education, race, ethnicity, and gender), caregiver-report questionnaires, and direct in-home assessments. These measures spanned multiple developmental domains, including language, cognition, motor development, social-emotional development, daily routines, developmental milestones, temperament, maternal emotional responses, parental perceptions of infant crying, maternal responsiveness, child behavior and parental stress, visual attention shifting, predictive looking, structured language production, language skills, gender socialization (via free play and preference surveys), and systemic influences of developmental milestones on family behaviors. Extended data Table 1 summarizes instruments and assessment timing. Parents completed questionnaires online, and in-home behavioral assessments were conducted when children were 1, 2, and 3 years old.

### Caregiver-Reported Questionnaire Data

The primary caregiver-report measure used in our analyses was the MacArthur–Bates Communicative Development Inventories (MB-CDIs^26^). Caregivers completed MB-CDI reports monthly beginning at 8 months of age and continuing through each child’s participation. We administered the Words & Gestures form from 8-16 months and the Words & Sentences form thereafter. Between 8 and 16 months, children’s language comprehension was additionally assessed through an interview-based administration of the Words & Gestures form.

In addition to language, caregiver-report measures captured social-emotional development^53,54^, developmental milestones, home routines, and broader family- and system-level influences on development.

### In-home experimental assessments

We also conducted a series of in-home assessments to characterize children’s developing cognitive and linguistic abilities. At years 1 and 2, children completed a home-based looking-while-listening task (online comprehension^55,56^), the Gap Overlap task (attentional shifting^56^), and the Pinwheel task (predictive looking^56,57^).

At approximately 24 months, children completed the Bayley-4 Scales of Infant and Toddler Development^58^, which provided standardized assessments of cognitive, motor, and language skills. At age 3 years, we administered the Quick Interactive Language Screener (QUILS^48^), a tablet-based assessment of vocabulary, syntax, and word learning. We also administered a study-developed book-reading task and tasks assessing gender perception^59–61^.

Together, these measures provide a multidimensional longitudinal profile of development across the first three years of life.

### Automatic data collection

#### In-home recording setup

The in-home system was designed not only for technical reliability, but also to minimize intrusiveness in everyday family life and to preserve family control over recording boundaries, including periods designated as explicitly private. We deliberately did not record overnight, even though caregiver–child interactions at night are very meaningful. Cameras turned off automatically after dinner. Families could decide exactly when and where to record. They could pause, relocate, or shut the system off entirely, whenever they wanted.

We used Wi-Fi cameras and far-field microphones to enable reliable, continuous in-home recording (camera: AXIS M1004-W Network Indoor Wireless IP camera; microphone: EveryWord© ultra far-field voice detection microphone). Audio was sampled at 16KHz PCM. Video frame resolution was 1920x1080 and sample rate 12 frames per second. Main living areas in each home (e.g., family room, playroom, kitchen, dining room) were equipped with one to two cameras with embedded microphones, along with one additional high-quality far-field microphone (see Figure 1). All devices were time-synchronized.

Hardware deployment and ongoing maintenance were conducted in partnership with Vector Security. Dedicated internet service was provisioned by Verizon Fios and fully segregated from families’ personal networks to support continuous upload. Because the system generated large volumes of data that had to be uploaded continuously, it required high upstream bandwidth (i.e., upload speeds comparable to download speeds), which is typically not available with standard consumer internet plans. Recordings were first routed to an intermediate secure server managed by EagleEye Security Management and then transferred to the AWS research environment.

#### Data download system

The system for downloading raw camera footage from temporary storage into the 1kD cloud environment comprised multiple modules implemented as AWS Lambda functions (Extended Data Fig. 1). These modules queried the temporary storage to identify newly available data and managed transfer into 1kD storage. Communication between modules was coordinated via AWS Simple Queue Service (SQS), enabling asynchronous, message-based exchange.

The download pipeline ran continuously. An hourly ***Enqueue*** job seeded the workflow by scanning all active devices and creating one retrieval task for each eligible hour slot. To improve reliability, tasks were lagged by 24 hours (i.e., the system requested data for the previous day’s hour blocks), allowing time for camera uploads to complete. Each retrieval message included the device identifier and a one-hour time range (start and end timestamps) and was passed to the *Retrieve* module. *Enqueue* respected each home’s recording schedule (e.g., 07:00–19:00 or a custom window) and generated tasks only for scheduled hours.

The ***Retrieve*** worker consumed task messages one at a time. For each device/time-range pair, it called the temporary-storage management API to list available recordings, persisted the listing and associated metadata in the 1kD database, and generated a footage-download message for each recording discovered. This design decoupled “discovery” from “download,” producing fine-grained work units downstream.

The ***Download*** worker consumed per-recording messages, requested the media from temporary storage, and wrote it into a two-week cooling storage tier. On success, it updated the recording’s state to “downloaded” in the 1kD database. Cooling storage served as a reversible buffer: families could request deletion during this window before data moved into long-term storage and processing.

Robustness and completeness were ensured via two ***Retry*** modules (*Retrieve Retry* and *Download Retry*). These modules scanned the 1kD database for hours or recordings where metadata retrieval or footage download had not completed successfully. When gaps were detected, they generated new messages to re-initiate the relevant steps, ensuring that missing data were eventually recovered and stored.

Scalability was achieved by operating at the finest practical granularity: messages were generated per device and per recording, on an hourly basis. This structure enabled concurrent processing of metadata listings and downloads across many workers, parallelization across devices and time blocks, and flexible scaling of compute resources as demand fluctuated.

#### The footage chunking and storage system

The system for chunking and storing footage consisted of multiple modules implemented using AWS Lambda services and EC2 instances (Extended Data Fig. 2). Modules were responsible for (1) transferring data from cooling storage to its permanent destination (raw data storage), (2) standardizing recordings by dividing them into one-minute chunks and writing them to the 1kD data lake, and (3) replicating storage to enhance resilience.

A daily ***file-sort*** Lambda moved data older than two weeks from cooling storage to raw data storage, where objects were locked using the AWS Object Lock mechanism. During this transfer, content was also replicated to a geographically separate AWS region to provide off-region backup.

Raw recordings varied in duration (from a few seconds to ∼5 minutes) and lacked consistent start/end boundaries. To standardize the corpus for downstream processing, we used a ***chunker*** module that processed 15-minute batches of raw footage and produced fifteen one-minute chunks aligned to whole-minute boundaries. A ***chunker scheduler*** regularly assembled and dispatched these 15-minute batches from raw storage. The resulting chunks were written to the 1kD data lake, protected with AWS Object Lock, and replicated for redundancy. Metadata associated with each chunk was logged to the 1kD MySQL database.

As in the download system, robustness and completeness were supported by a *scheduler-retry module that* identified batches for which chunking had not completed successfully and re-queued them for processing. Scalability was achieved through AWS auto-scaling: the number of chunker instances was adjusted based on the current volume of footage awaiting chunking. Data resilience was maintained through object locking and cross-region replication.

### The scalable data analysis pipeline

#### The batch processing system

The batch processing system comprised several AWS-based modules implemented using AWS Glue, ECS, and DynamoDB (Extended Data Fig. 3). At its core was a DynamoDB table - the 1kD feature table - that stores all available annotations at any given time (e.g., speech detected per room, number of people present). The feature table was integrated with the 1kD metadata database so that the full inventory of available footage was represented alongside derived annotations.

Algorithms were executed on individual files or defined subsets via an Algorithm Runner and associated model Docker containers, which exposed model APIs. A Results Collector module then gathered raw model outputs, normalized them, and applied additional logic to produce consistently structured annotations suitable for downstream scientific analyses.

The ***Algorithm Runner*** was the core component of the system. It was implemented as an Apache Spark job running as an AWS Glue job within the secure cloud environment and managed the end-to-end annotation process - from selecting inputs to persisting outputs. Specifically, it accessed the table of collected segments, recorded minutes, and applied a predefined configuration to select subsets for analysis. Subsetting could be based on date range, household, and outputs from prior algorithms, among other criteria. Once selected, the runner launched a distributed workflow: basic processing units (individual files or small groups of files) were dispatched as HTTP requests to pre-instantiated Docker containers hosting the relevant algorithms. Results were returned to the runner, written initially to S3, and then ingested into the final database via a separate Glue job.

Algorithms varied in complexity, from lightweight operations (e.g., audio or frame extraction) to more computationally intensive models (e.g., transcription). Each algorithm was containerized and deployed on EC2 instances (CPU or GPU) within an ECS cluster. To accommodate fluctuating demand, AWS auto-scaling provisions instances dynamically as needed. To improve cost efficiency, we used AWS spot instances when feasible; although availability was not guaranteed, they substantially reduced compute costs relative to on-demand instances.

The ***Results Collector*** served as the second major component of batch processing, aggregating and transforming raw model outputs into analysis-ready annotations. Collector operations ranged from simple transformations (e.g., converting speech duration to a binary speech presence label) to multi-model aggregation. For example, a coarse-aggregation collector combined motion- and speech-detection outputs to assign a binary label to each recorded minute, indicating whether it should be excluded from further processing (e.g., no one was present at home) or retained for subsequent analysis.

Overall, the batch processing system was designed to be flexible and cost-efficient. Across runs, we executed approximately eight core algorithms (audio/frame extraction, motion detection, speech detection, speech enhancement, transcription, and several collectors) over batches ranging from ∼10^5 to 5×10^6 files. Depending on batch size, model mix, and CPU/GPU configuration, processing times ranged from ∼1.5 hours to ∼3 days. Costs were driven primarily by model runtime and hardware type; where possible, we parallelized within instances to improve throughput per dollar.

#### Batch processing using Large Language Models

We inferred who was present, minute by minute, using a large multimodal language model (LLM) trained on a single representative frame per video (the mid-minute frame). Restricting inference to a single frame per minute per camera substantially reduced API/compute cost while preserving a consistent snapshot for presence detection. Frames were drawn only from minutes pre-flagged as “likely occupied” by lightweight speech and motion-detection.

To operate at scale, we used the provider’s batch API: images were sent in batches of 200 with ∼60 concurrent batch requests. A controller service queued batches, monitored their status, and downloaded completed results; failures were retried with back-off. All outputs (and any errors) were parsed into a schema-validated results table. All annotations were completed within a secure, institutionally managed computing environment, with access restricted to authorized personnel and in accordance with our approved data-security protocols.

The prompt requested yes/no judgments per frame for: (i) infant presence (with an age threshold that shifted over time—“under 2 years” early, “over 2” later), (ii) adult female, (iii) adult male, and (iv) another child older than ∼2–3 years (age-appropriate threshold). The volume of annotated frames per family scaled with time at home and device count; as an order of magnitude, annotating ∼1M frames required ∼24 hours end-to-end.

### Algorithms deployed in the pipeline

We describe the implementation details of the algorithms deployed in the analysis pipeline (Fig. 4): motion detection, speech detection, infant-presence detection, transcription (including recording selection), and cross-modal aggregation and integration. With the exception of transcription and the final aggregation stages, all modules operated on individual audio snippets or single video frames.

### Motion detection

#### Objective

Identify the minutes of recordings that show potential human activity in each room and distinguish them from the times when the room is empty. This allowed us to filter the data and focus our analysis on behaviorally informative segments.

#### Rationale

Human presence typically entails movement. While this heuristic is not perfect, as it also flags pet motion or lighting fluctuations, it allowed us to remove minutes in which we were highly confident that no interesting behavior was taking place. Sleep immobility is not a primary concern in our current work. Such instances are filtered out and reserved for future work.

#### Method

For each video, we compared temporally adjacent frames sampled every *x* seconds. We computed frame differences and threshold them at value *th*; connected components (“contours”) above *th* were extracted, and a clip was labeled **motion** if any contour exceeded an area threshold *A*. Thus, a minute was flagged when it exhibited sufficiently large, sustained pixel changes.

### Speech detection

#### Objective

Identify minutes that contain speech so that heavy downstream processing (e.g., transcription) can focus on linguistically informative segments and skip silence.

#### Rationale

A high-recall sound detector was used to eliminate silent periods and capture all recorded moments deemed to contain speech.

#### Method

For each 1-minute audio segment, we slid fixed-length, non-overlapping 4 s windows. Each window was passed to a Wav2Vec-based transcriber^32^. A window was tagged as “speech” if any meaningful transcription was produced; the segment’s speech length was the count of speech windows × 4 s, and the minute was flagged as “speech present” if speech length was greater than 0. The 4-second window balanced two factors: a shorter window length enhanced precision in determining the total number of seconds during which speech was perceived within the household for a given recording; however, below 4 seconds, accuracy could be compromised, as transcription required sufficient context.

### Speech transcription

#### Objective

Transcribe speech occurring within a given house area (room or open space).

#### Rationale

Each area typically has multiple simultaneous recordings (e.g., overlapping mics/cameras), creating redundancy. To maximize transcript quality and control cost, we first selected the highest-quality recording per minute for that area, then transcribed only that signal.

### Method

#### Signal selection via speech energy

For every candidate audio recording in a minute, we estimated a speech-focused energy score used as a proxy for transcription quality. We first denoised the waveform with a neural speech enhancer^33^, which suppressed background noise while preserving speech. We then computed the aggregate energy of the enhanced signal (sum of absolute amplitudes for the full minute). The recording with the highest enhanced-speech energy was selected for transcription.

#### Transcription

The selected audio was transcribed with a state-of-the-art ASR model (WhisperX^34^). We processed 30-second chunks (the model’s practical limit), using WhisperX large-v2 with temperature = 0, VAD onset = 0.20, and VAD offset = 0.363. To make downstream comparisons fair, we applied a text normalization pass to transcripts: removed non-lexical vocalizations (e.g., “oh”, “ah”, “mhm”), striped punctuation, and applied automatic spelling normalization.

### Baby detection

#### Objective

Determine whether an infant is present in a given area of the home.

#### Rationale

Over the course of the three-year study, the infant grew, changed, and their behavior and appearance evolved accordingly. Each area of the home was covered by different recording devices with varying fields of view, and people - especially infants - moved frequently through space. As a result, the infant’s location at any given moment could be difficult to detect from a single viewpoint. To ensure accurate coverage, we required broad annotation across multiple household areas for each candidate minute.

#### Method

For every minute in which we suspected that the baby (or any other person) was in the home, all available devices and cameras were considered. Because annotating multiple frames per minute was expensive, we annotated only a single frame per minute: the middle frame (at 30 seconds). This frame was analyzed using GPT-4o (within a secure, enterprise-grade environment configured for research use) with the following prompt: “Please answer yes or no to each question, with each response in a new line. Try your best even if you are not sure. (1) female adult, (2) male adult, (3) baby younger than 2 years, (4) child older than 2 years (sibling).” Ages were adapted based on the exact date annotated and the family composition. The model’s responses were parsed into binary indicators (1 or 0) and stored in a database. No parameter tuning was performed.

### Cross-modal integration and aggregation

#### Objective

Isolate minutes in which the infant is present and pair them with the speech in the infant’s vicinity.

#### Rationale

The combination of multiple modalities, temporal variability, device redundancy, and naturalistic noise necessitated aggregation across space and time. To remain cost-effective, we first applied lightweight detectors to flag candidate minutes, then ran heavier analyses (baby-presence detection and transcription) only on those minutes, and finally applied temporal smoothing before making a minute-level decision. All devices in the setup were time synchronized (see in-home recording setup).

### Method

#### Spatial layout and device–area mapping

Each home was partitioned into non-overlapping vision areas (rooms or contiguous open spaces). Every camera that “looked into” an area - whether physically located in that area or viewing it from an adjacent room- was mapped to it. In parallel, we defined non-overlapping speech areas by grouping microphones that captured audio from a given physical space; each speech area was linked to a vision area.

### Per-signal aggregation (per minute, per area)

#### Motion (vision)

We averaged motion scores from all cameras mapped to the area to obtain a single area-level motion score.

#### Baby presence (vision)

We computed a weighted average of infant-presence scores, assigning a weight of 1.0 to cameras located within the area and 0.5 to cameras outside the area that still viewed it; this favored primary viewpoints while retaining evidence from auxiliary angles.

#### Speech (audio)

For each speech area, we chose the maximum detected speech duration across its devices, assuming that the best-placed microphone captured the longest signal.

#### Activity gating and annotation

A minute was flagged as active if the home showed both (i) motion > 0 and (ii) speech > 0 anywhere, or (iii) any area had ≥ 8 s of detected speech. Only active minutes were used for infant-presence annotation. For transcription, we processed every minute with any detected speech (> 0 s).

#### Temporal smoothing

To stabilize infant-presence evidence, we smoothed each area’s weighted presence score using a symmetric 5-minute window (past and future). Specifically, we applied decaying weights (0.75, 0.50, 0.25, 0.10) for ±1 to ±4 minutes around the processed minute.

#### Minute-level location and transcript assignment

For each minute, we (1) thresholded smoothed presence scores to suppress low-confidence detections; (2) selected the infant’s location as the area with the maximum remaining score; and (3) attached the transcript from the speech area linked to that vision area as the infant’s proximal linguistic input.

#### Summary

The algorithmic processing pipeline fused redundant multi-device signals, restricted expensive processing to behaviorally informative segments, exploited the temporal structure of the data, and produced a minute-resolved assignment of infant location with aligned speech content.

### Corpora language analysis

#### Dictionary creation

For the household recordings, we used the Child+Speech split, with the child-presence threshold set to 0.1 (Table 1). For CHILDES, we used the version distributed through the BabyLM Challenge^62^.

All text was processed with spaCy^63^. Transcripts were first segmented into sentences and then tokenized. Each token was annotated with its original surface form, lemma, and part-of-speech (POS) tag. We then applied a standardized normalization and filtering pipeline to reduce spurious lexical variation and transcription artifacts. To ensure proper nouns contributed to noun analyses, we reassigned PROPN tags to the noun category (PROPN → NOUN) rather than treating proper nouns as a separate class. Surface forms and lemmas were lowercased and filtered to retain only word-like items. Specifically, we retained only tokens whose lemmas contained alphabetic characters, excluded items containing digits, and removed punctuation and whitespace tokens. To further reduce noise, we removed single-letter tokens except for the English words “a” and “i.” We also denoised lemmas by collapsing character elongations, reducing runs of more than three identical characters to two. After normalization, analyses were either performed on the full dictionary or restricted to a POS subset (e.g., nouns, verbs).

Vocabulary frequencies were computed by grouping tokens by lemma and counting occurrences, yielding a unigram frequency distribution over lemmas and the corresponding total token count for each corpus.

#### Reference corpora and cross-corpus comparison

To quantify cross-household similarity in lexical usage, we compared each family corpus to an explicit reference corpus whose unigram distribution served as the baseline. We used CHILDES and, in turn, each of the eight processed family corpora as the reference; for each reference, the comparison set consisted of the remaining 1kD households corpora. For each word type in the reference vocabulary, we asked how many comparison corpora showed a frequency consistent with the reference. Let 𝐶*_i_* (𝑤) be the count of word type w in comparison corpus i, and 𝑁i the total token count in that corpus. To stabilize estimates for low-frequency types and avoid undefined ratios, we estimated probabilities with additive smoothing: 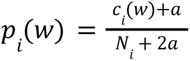, where 𝑎 = 0. 5. Reference probabilities 𝑝_ref_(𝑤) were computed analogously. For each corpus i and word w, deviation from the reference probability (𝑝_ref_(𝑤)) was quantified as a symmetric ratio: 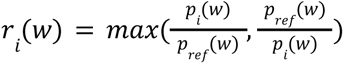. This measure is always at least 1, equals 1 when corpus i matches the reference for w, and increases as w becomes relatively over- or under-represented. A word type was considered shared between corpus i and the reference if 𝑟*_i_*(𝑤) ≤ 2, that is, if its estimated probability in the comparison corpus fell within a twofold band of its probability in the reference corpus. For each reference corpus, we then counted, for each word in the reference vocabulary, the number of comparison corpora that met this criterion. We also explored a frequency-dependent statistical tolerance that allowed slightly wider deviations for low-count words, but this changed shared versus not-shared classifications only minimally and did not materially affect the overall results; we therefore report the simpler fixed twofold criterion.

## Supporting information

Full Supplementry

## Extended data

**Extended data table 1.** Summary of behavioral measurements collected during the study. . including domain, instrument, and assessment timing. Measures include caregiver-report questionnaires, behavioral tasks, qualitative interviews, and standardized developmental assessments collected at repeated and age-specific time points.

| Domain | Instrument | Timing |
| --- | --- | --- |
| Language Comprehension and Production | MacArthur-Bates Communicative Development Inventories (MB-CDIs<sup>26</sup>) | Monthly from the age of 8 months |
| Routines | Home Routines Questionnaire (Study developed) | 12 months |
| Developmental Milestones: fine and gross motor skills, language, social interactions, and cognitive development | TUFTS Well-Being of Children Scale<sup>64</sup>. | 23–28 months |
| Temperament | Early Childhood Behavior Questionnaire<sup>54</sup> | 20 months |
| Crying Behavior | Mothers' Emotional Responses to Infant Crying<sup>65</sup>, 2021), Infant Crying Questionnaire<sup>53</sup>, Maternal Responsiveness Questionnaire<sup>66</sup> | 6 months |
| Parental Stress | The Parenting Stress Index (PSI-4<sup>67</sup>) | 28 - 30 months |
| Systemic influences of developmental milestones—such as motor skills, sleep, breastfeeding, and social development—on family behaviors and interactions. | Qualitative Interview (Study developed) | 33 months |
| Gender Development | Behavioral Assessment<sup>59–61</sup> | Year 3 home visit |
| Attention and language processing | Looking While Listening (online comprehension<sup>55,56</sup>), the Gap Overlap task (attentional shifting<sup>56</sup>), and the Pinwheel task (predictive looking<sup>56,57</sup>) | Year 1 and 2 home visit |
| Language | Quick Interactive Language Screener (QUILS<sup>48</sup>) | Year 3 Home visit |
| Language production | Book Reading Task (Study developed) | Year 3 Home Visit |
| Language Cognitive and Motor development | Bayley's Scales of Development<sup>58</sup> | End of second year |

**Extended Data Fig. 1.**
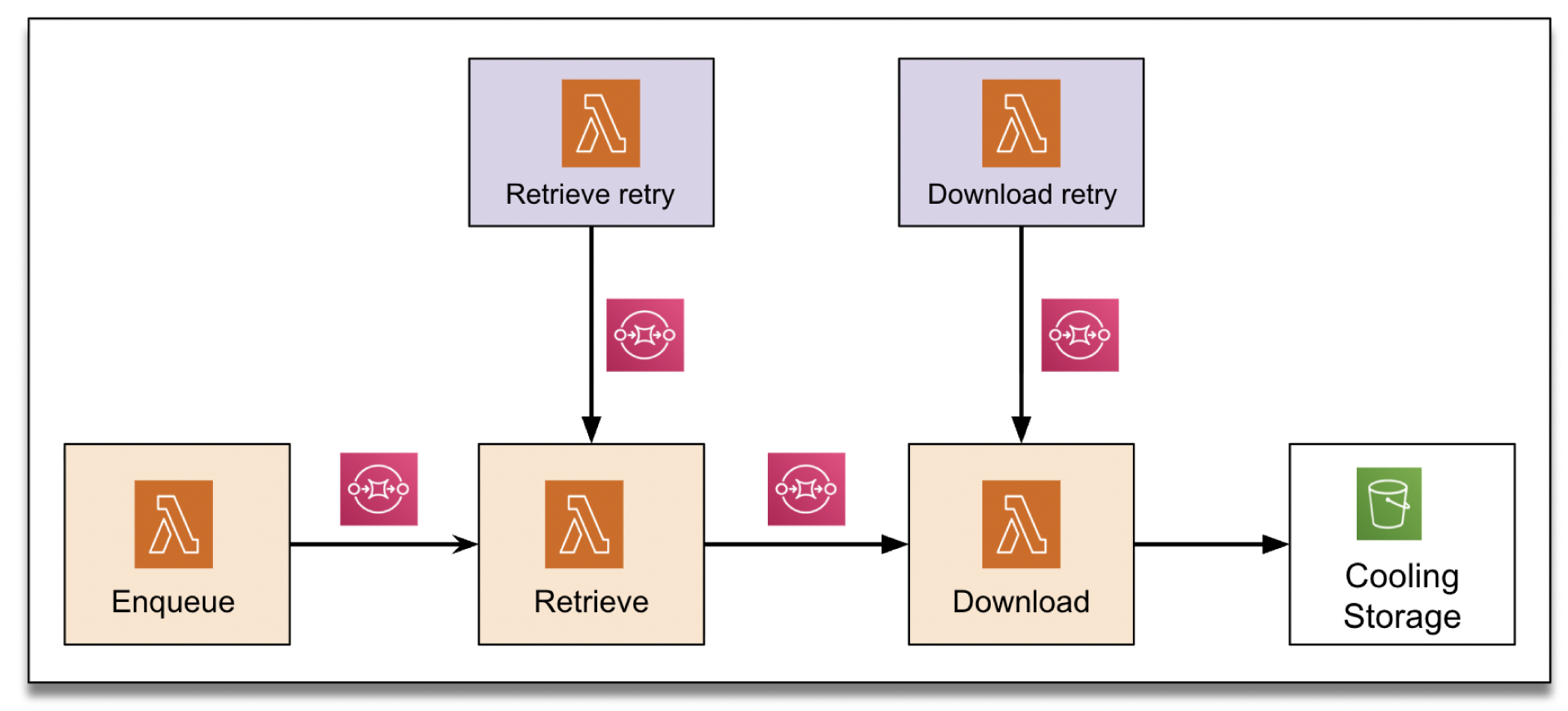
Architecture of the 1kD data download system. An hourly Enqueue job created retrieval tasks for each eligible (device, hour) slot; the Retrieve module discovered available recordings and documented their metadata, and the Download module fetched each recording into a two-week cooling storage tier. Retry services monitored the 1kD database for gaps and regenerated tasks when needed, ensuring robust, hour-level ingestion of home camera footage into the long-term processing environment.

**Extended data Figure 2.**
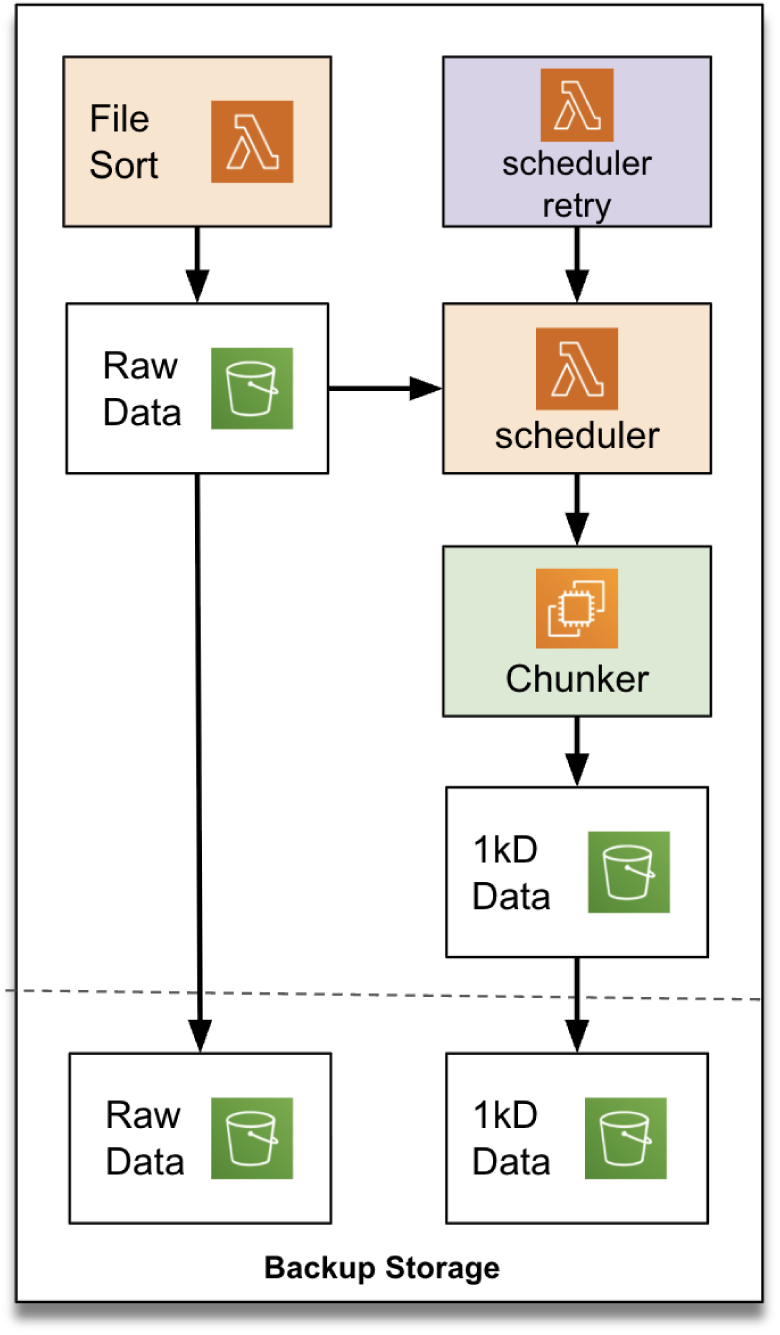
Footage chunking and storage system. Data older than two weeks were moved from cooling storage to locked, replicated raw storage by the File Sort job. A Chunker Scheduler assembled 15-minute batches of raw footage, which Chunker workers converted into one-minute, minute-aligned chunks stored in the 1kD data lake with backup replication. A scheduler retry component and AWS auto-scaling together provided robust, scalable processing.

**Extended Data Fig. 3.**
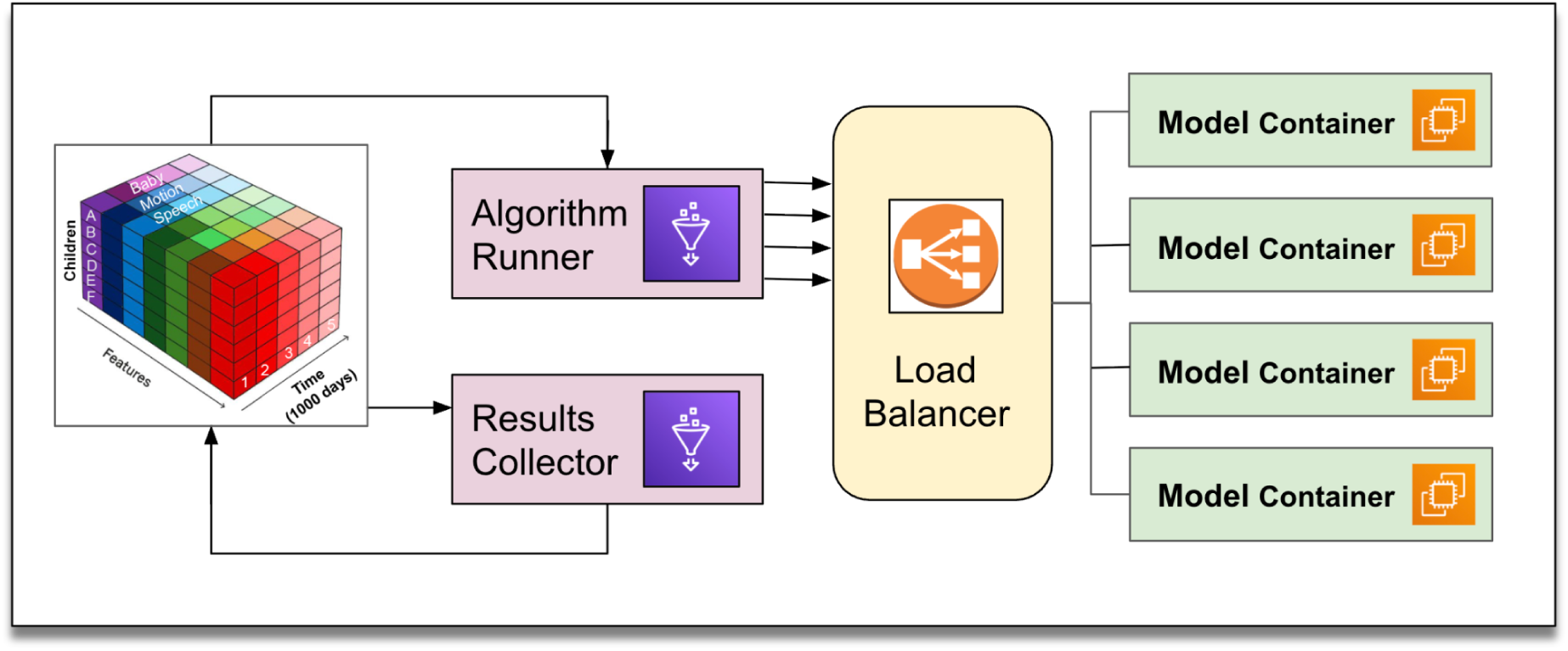
Batch processing system. The algorithm runner (an AWS Glue job) queried the 1kD feature table in DynamoDB to select subsets of footage and sent requests to models containers running on EC2/ECS. Model outputs were written to the feature table. Collector jobs were used to normalize and aggregate results.

**Extended data Figure 4.**
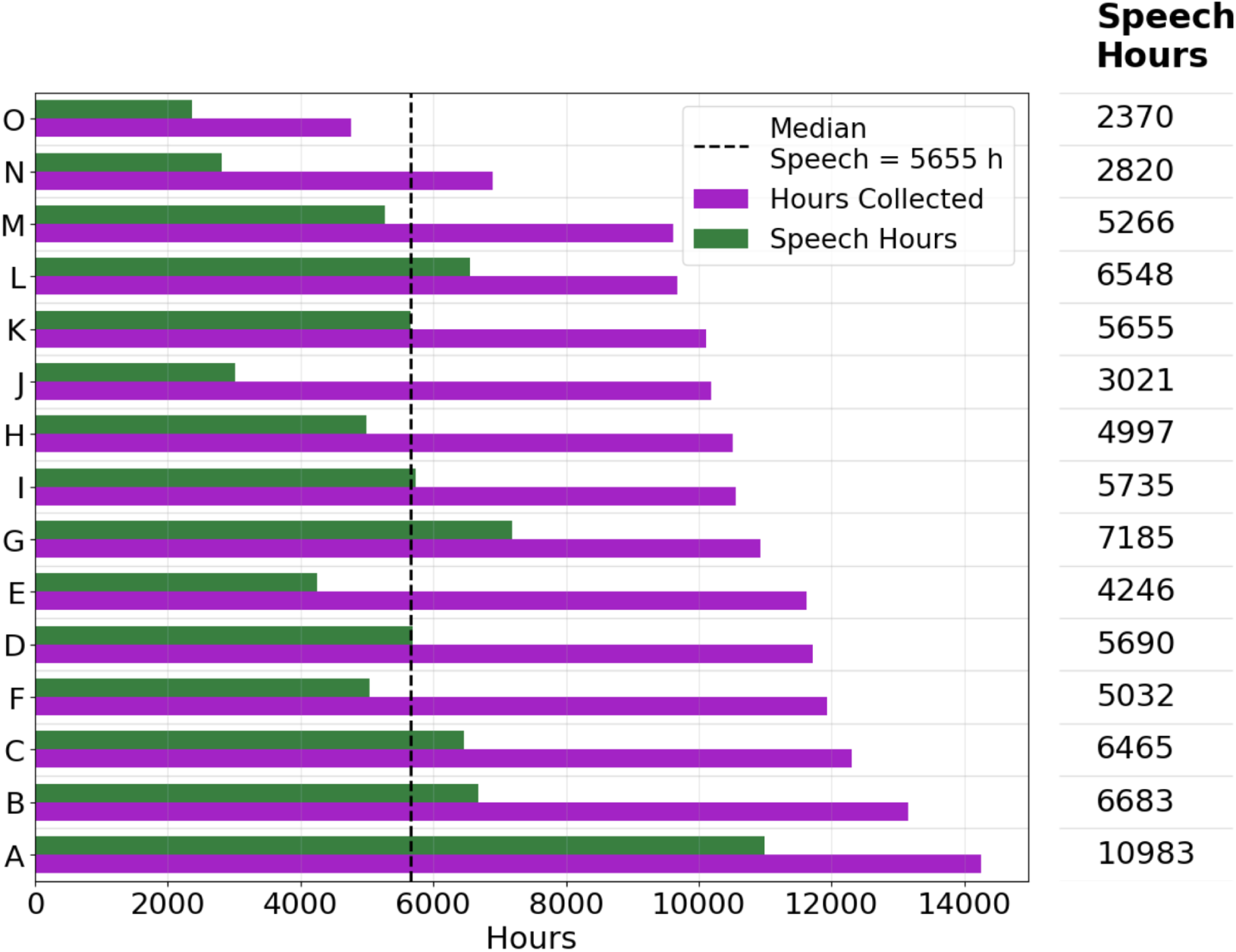
Total recorded hours and isolated speech hours per family. Bars show, for each household, the total unique recording hours (purple) and the subset of hours in which speech was detected by the audio pipeline (green, range: 2,370–10,983 speech hours; median: 5,655). Homes are sorted by the total number of hours collected.

## Acknowledgements and funding

We extend our heartfelt gratitude to the 1kD families for their remarkable contributions, generosity, and invaluable cooperation throughout this project. At Princeton, we would like to express our sincere appreciation to Irene Kopaliani, John Wiggins, Paula Looney, Robert Sullivan, Elizabeth Adams, Mark Ratliff, and the Princeton Institutional Review Board, along with many others across the university, whose exceptional institutional support played a crucial role in making this unconventional project a reality. We are also deeply thankful for our collaboration with our partners at Sigma Software, especially Stanislav Samko and Olga Chikhiro, whose dedication and thoughtfulness were pivotal in providing the technical expertise needed to build the software infrastructure for this initiative. Additionally, we wish to acknowledge Syed Ahmad of Beaty Consultancy for his contributions. Our sincere gratitude extends to the Vector Security team in Princeton, New Jersey, as well as Eagle Eye Networks and ArkX Labs, for their outstanding partnership and expertise in developing the necessary hardware infrastructure that has made this project possible. We are also grateful to Sergei Drobner, Eric Co, Alex Christodoulou, Matthew Funcke, and Samantha Pramanick for their contributions to the design and development of the data collection pipeline. We thank Natalie White, Ore Segal, Idan Margalit, Ajay Donthula, Asa Santos, Siniru Iehoma, Eric Kidder, Aarushi Adlakha, and Sara Lougy for their contributions to data annotation and project support. We thank Holly Baines, the Program Director at Wellcome Leap, for her invaluable initial support. This work was supported by Wellcome Leap, the McGregor Girand Charitable Endowment, and several grants from Princeton University, including Princeton Language and Intelligence, Princeton Precision Health, Data-Driven Social Science, and the AI Lab. The content of this publication and any related materials is solely the responsibility of the authors and does not necessarily represent the official views of the funding agencies.

