## Supplementary material for "The First 1,000 Days (1kD) Project - Collecting and Analyzing an Ultra-Dense Naturalistic Dataset of Human Baby Development": Full Supplementry

The 1kD dataset captures everyday life as it naturally unfolds - without scripts, staging, or observer interference. This results in high variability across time, space, and social context, reflecting the dynamic and often unpredictable nature of real home environments. The figures below provide illustrative examples of this variability, highlighting both spatial and auditory complexity across different families and time windows.

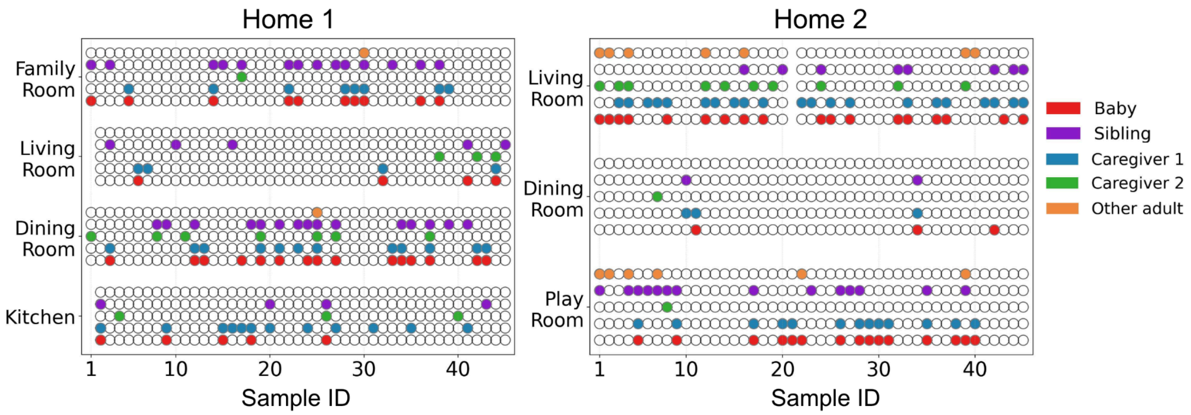


**Supplementary Figure 1. Spatial presence across household areas.** Spatial presence of individuals across sampled time segments for two households. Each vertical line represents a single time sample. Colors indicate individual identities: red = baby, purple = sibling, blue = caregiver 1, green = caregiver 2, and orange = other adult. Samples are non-consecutive and not taken from the same day.

Supplementary Figure 1 presents the spatial distribution of individuals across household locations for two families, based on sampled one minute recordings. Importantly, the samples are not consecutive and are not drawn from the same day. Instead, they reflect a set of diverse time windows selected across the data collection period. The figure illustrates clear variability in where people are located, with different configurations of individuals co-present in different household areas. This variation highlights the dynamic nature of daily interactions within the home environment, and the importance of capturing variability at scale for understanding and modelling the complexity of real-world, in-context, developmental processes.

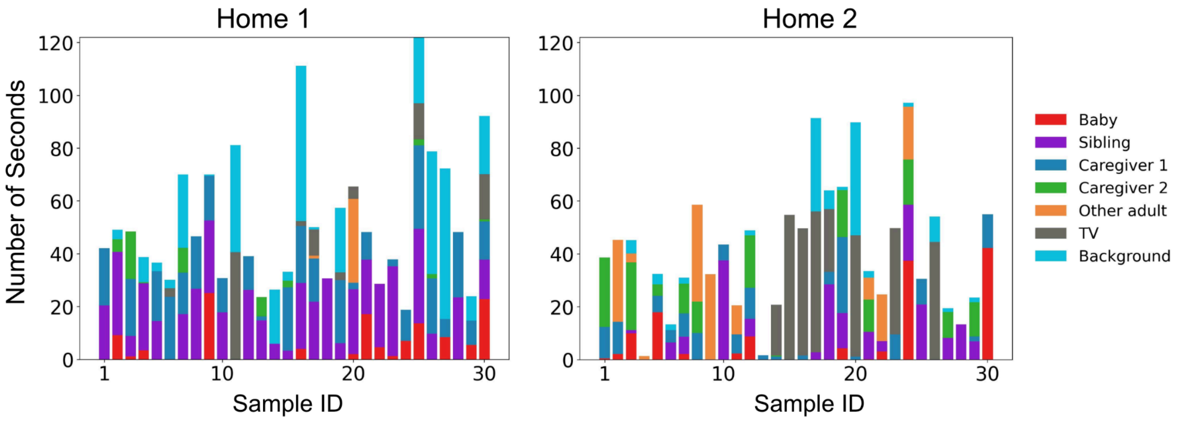


**Supplementary Figure 2. Speech type distribution within households.** Duration of speech-related sounds per minute, showing overlapping and concurrent sources in the home environment. Color codes represent different sound types: Red = baby vocalizations, Purple = sibling speech, Blue = caregiver 1 speech, Green = caregiver 2 speech, Orange = other adult speech, Grey = television, Light blue = background household noise (e.g., music, laughter, mechanical sounds).

Supplementary Figure 2 presents the total number of seconds attributed to various speech-related audio categories within one-minute intervals for two families. These categories include adult speech (from caregivers or other adults), baby and child vocalizations, television sound, and background noise (e.g., household sounds). Notably, the total often exceeds 60 seconds per minute, as overlapping events are plotted at their full duration even when they occur simultaneously. The samples were randomly selected across the data collection period.

This visualization reflects the rich auditory complexity of the home environment, where multiple sound sources frequently co-occur. In addition to primary vocalizations, the presence of background noise, television, and overlapping voices contributes to a layered soundscape - posing significant challenges not only for automated detection systems but also for developing listeners, such as the babies themselves.

### Evaluation of analysis-pipeline algorithms

End-to-end performance of the full pipeline is reported in the main paper. Here, we report per-algorithm metrics computed on sampled subsets of families collected over the course of the study. Although these values are not highly informative in isolation, we include them to provide a sense of individual algorithm performance. Full model descriptions and parameter settings are provided in the Methods section.

**Motion detection.** For each household, 100–250 one-minute clips were hand-labeled as containing motion or no motion. We selected model parameters (*th*, *A*) to maximize F1 on each household's validation set, searching $th \in \{5, 20, 30, 40\}$ and $A \in \{0, 100, 200\}$. This per-home calibration accommodated differences in layout, lighting, and device placement. The average F1 across homes was 0.89. Mean recall was 0.88. The motion signal was aggregated across devices and combined with other cues (e.g., speech) in the end-to-end pipeline.

**Speech detection.** For each of 9 families, we hand-labeled 100–250 one-minute clips. We computed average precision and recall separately for speech-present and silence minutes. Silence precision and recall were 0.93 and 0.83, respectively, whereas speech precision and recall were 0.83 and 0.92, respectively. As intended for this filtering stage, silence detection achieved high precision (few false speech calls in quiet minutes), whereas speech detection achieved high recall (few misses when speech was present). Performance improved as the duration of speech within a minute increased, with very brief or isolated utterances being harder to detect. No per-home hyperparameter search was required beyond fixing the 4-s window, and this setting generalized well across households.

**Speech transcription.** No tuning was performed. The evaluation measures results are described in the Results section.

**Baby detection.** We evaluated single-frame annotation quality on 398 frames sampled from four homes, including 80 frames with infant presence. Baby detection achieved a precision of 0.71, recall of 0.91, and F1 of 0.80. Given the low base rate of infant presence, the observed precision was more than 3.5-fold above chance. For other individuals, sibling detection achieved 0.83 precision and 0.97 recall (30 occurrences), female adult detection achieved 0.88 precision and 0.85 recall (124 occurrences), and male adult detection achieved 0.70 precision and 0.93 recall (55 occurrences). End-to-end evaluation across minutes and households is reported in the Results section.

**Cross-modal integration and aggregation.** Quantitative performance metrics are reported in the Results section. Supplementary Fig. 3 presents F1 scores for Infant+Speech detection (A) and (Any) Child+Speech detection (B) as a function of the infant or any-child presence detection threshold. As expected, F1 declined as the threshold increased, owing to lower recall despite higher precision. Performance nevertheless remained high at lower thresholds and often exceeded 0.8 for most families. Higher thresholds selected instances with a very high probability of infant presence, but reduced overall coverage. Each colored line corresponds to a single family. Detection performance varied across households, likely reflecting differences in home layout, household composition, and camera visibility.

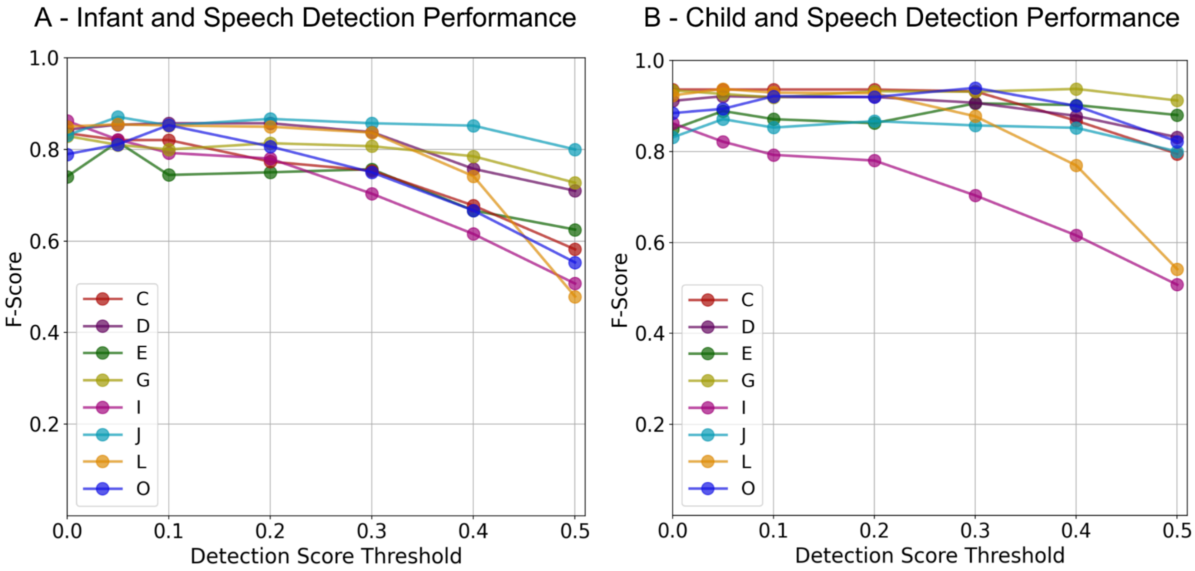


**Supplementary Fig. 3. End-to-end pipeline performance.** F1 score for speech detection in the presence of a child. Each line represents a different family. (A) F1 score for infant+speech detection across infant-detection score thresholds. (B) F1 score for any child+speech detection across detection score thresholds.

### Transcription annotations guidelines

Annotators were provided with all video and audio recordings from a specific minute. Their task was to transcribe the highest-quality recorded speech, capturing both the content of the speech and identifying the speakers. Annotations were made by marking segments within the selected audio. More specifically they were provided with the following guidelines:

<u>Speech annotations:</u>

- An utterance is defined as a natural unit of speech bounded by silence or a change in speaker. Speech segments should typically contain one or up to a very few utterances.
- A speech segment should be labeled with the identity of the speaker when possible.
    - Identifiable speaker: Labels like **"father"** or **"mother"** should be used if the speaker is recognized.
    - The **"crosstalk"** label should be used when more than one person is talking simultaneously.
    - The **"other person"** label should be used when an unidentifiable person is talking.
    - Speech from a TV or another electronic device should be labeled **"tv adult"** or **"tv child"** depending on if the speaker is an adult or child.
    - Singing or music with lyrics should be labeled as **"music".**

Non-speech sound labels:

- Sounds of the following categories should be marked with a segment and labeled using the category name: noise, laughter, dial tone, baby crying, baby sounds, child crying, child sounds, or faint (used for indistinct or unidentifiable speech).
- If sounds do not co-occur with speech they should be segmented separately. Otherwise, the speech segment should be added with the relevant label.
- Noises like coughing, sneezing, or yawning should not be segmented or annotated.

Content transcription:

- **Utterances should be transcribed as is**—including stops, restarts, vocal fillers (“like”, “um”, “uh”, “erm,” etc.), stuttering, and both speech and grammatical errors.
- Interrupted elements should be marked with a short dash (“-”)
    - Ex: “Well, it’s because-”
- Sound elongation should not be marked.
    - Ex: “Soooooo, the other day…” should be rendered “So, the other day…”
- Duplication or restarts should be marked using a short dash (“-”).
    - Ex: “I was- I was sitting”
- If the speech is unclear, the text [unintelligible] should be used exclusively for the unintelligible portion of the utterance.
- If the person is speaking a language other than english - [foreign] should be used for the portion of foreign language
    - Ex: “In spanish it’s uno dos tres” it should be written “In spanish it’s [foreign].”
- Interruptions or overlapping speech should be transcribed when possible, by separating the multiple speakers into separate segments. If there is no way to separate the speech into individual segments, a dominant speaker should be chosen and transcribed as normal while adding the interjections from the second speaker using square brackets
    - Ex: “Hello! How are you [I’m doing fine] today? Oh that’s good [Oh I didn’t mean to cut you off there.]”
- Any sound of active listening (“yeah”, “mmm hmm”, “okay”, etc.) does not need it’s own segment
    - Ex: “So I went up to him [Mhmm] and I told him the plan”
- Periods should be placed as a speaker naturally concludes a sentence.
- Comma should be used for short pauses. When a speaker trails off, an ellipsis should be used (“...”).
- Quotes, both direct (ex: the text said “see you at school”) and indirect (ex: I was like “I can’t believe he did that”), should be put in quotation marks.

- Proper practices for capitalization and spelling should be followed.
- A question mark should be used after questions and utterances intended as a question.
- Child speech should be transcribed with errors verbatim (errors should be included as spoken)
    - Ex: “I raned so fast I felled”
- Sounds of assent like “mmm” or “hmm” should be written as spoken.
- Any other communicative speech sound like “oh” or “ah” should be written as spoken.
- Abbreviations, acronyms, or shorthands should not be used unless the speaker uses them.
    - Ex: “I was in the US working for UPS”
    - Ex: if someone said “BT dubs” instead of “BTW” it should be transcribed verbatim as “BT dubs”
- Contractions should be transcribed as spoken.
    - Ex: “I have been really tired lately”
- Symbols should not be used to replace words that can be written verbatim.
    - Use Dollar or dollars instead of $
    - Use And instead of &
- Numbers should be written as plain english text
    - Ex: “one thirty in the afternoon”.

#### Linguistic properties of natural home environments - Part of Speech analysis

To test whether the full-dictionary sharedness pattern reported in the Results section held within major grammatical classes, we repeated the same analysis separately for verbs and for nouns. Results are shown in Supplementary Fig. 4- A (Verbs) and B (Nouns), with stacked bars indicating, for each reference corpus, the percentage of tested words that were matched (within the allowed ratio threshold; Methods) by 7–9 corpora, 5–6 corpora, 2–4 corpora, or by no other family (“unique”).

Overall, the distributions show both shared structure and systematic differences across lexical categories. Verbs were more broadly shared across households: for most reference corpora, a larger proportion of verb types fell into the highest sharedness bin (shared with 7–9 corpora), with a smaller proportion in the lowest sharedness bins. Nouns showed lower cross-corpora convergence, with more noun types concentrated in the intermediate and low sharedness bins (5–6 and 2–4 corpora) and a larger unique component, indicating greater household-to-household variability in noun frequencies. When CHILDES served as the reference, the unique fraction increased for both parts of speech, indicating that a larger set of reference vocabulary items lacked frequency matches in household language under the same tolerance criterion. Importantly, the relative pattern remained consistent across reference choices: verbs were generally more similar across corpora than nouns.

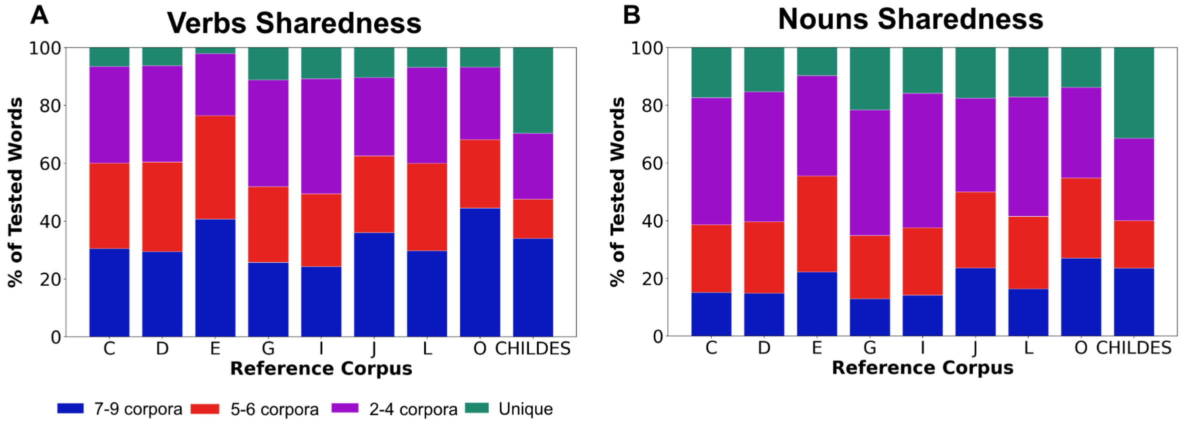


**Supplementary Figure 4. Verb and noun sharedness across households.** For each reference corpus (families C, D, E, G, I, J, L, O; and CHILDES), we assessed, for verbs (panel A) and nouns (panel B) how many other family corpora matched the reference frequency (Methods). Stacked bars show the proportion of tested vocabulary words that were shared with 7–9 corpora, 5–6 corpora, 2–4 corpora, or were unique (matched by none).
